# Dendritic domain-specific sampling of long-range axons shapes feedforward and feedback connectivity of L5 neurons

**DOI:** 10.1101/2021.01.31.429033

**Authors:** Alessandro R. Galloni, Zhiwen Ye, Ede Rancz

## Abstract

Feedforward and feedback pathways interact in specific dendritic domains to enable cognitive functions such as predictive processing and learning. Based on axonal projections, hierarchically lower areas are thought to form synapses primarily on dendrites in middle cortical layers, while higher-order areas are posited to target dendrites in layer 1 and in deep layers. However, the extent to which functional synapses form in regions of axo-dendritic overlap has not been extensively studied. Here, we use viral tracing in the secondary visual cortex of mice to map brain-wide inputs to thick-tufted layer 5 pyramidal neurons. Furthermore, we provide a comprehensive map of input locations through subcellular optogenetic circuit mapping. We show that input pathways target distinct dendritic domains with far greater specificity than appears from their axonal branching, often deviating substantially from the canonical patterns. Common assumptions regarding the dendrite-level interaction of feedforward and feedback inputs may thus need revisiting.

## Introduction

One of the key organizing principles of connectivity within the neocortex is thought to be hierarchy between cortical areas. This notion was originally proposed by Hubel & Wiesel (Hubel & Wiesel 1962) to account for the increasing receptive field complexity in areas progressively further from the retina. In purely feedforward (FF) networks, such as artificial neural networks used successfully in computer vision (LeCun et al 2015), hierarchy is generally defined by synaptic distance from the sensory periphery. However, in highly recurrent networks like the cortex, it is not possible to apply this definition consistently beyond the initial levels. Instead, the laminar patterns of axonal projections are often used to deduce relative levels of hierarchy. For example, the projection from primary to secondary visual cortex, which is classically defined as FF, is characterised by dense axon terminations in middle cortical layers (particularly L4). Meanwhile, the projection from secondary to primary visual cortex, used as the basis for defining feedback (FB), primarily targets L1 and to a lesser extent deeper layers (Rockland & Pandya 1979). This pattern of FF and FB projections also appears in many other brain regions, and has been used as a proxy to describe the hierarchical relationships between a large number of areas across the brain (D’Souza et al 2016, D’Souza et al 2020, Felleman & Van Essen 1991, Harris et al 2019, Wang et al 2020b, Zeng 2018).

Projections, however, do not guarantee functional connections. The link between the two is called Peters’ rule, which postulates that the probability of synaptic connections can be predicted from the overlap between axonal and dendritic arbours (Rees et al 2017). While overlap between axons and dendrites is necessary for synapses to form, it is far from sufficient. The link between axo-dendritic overlap and connection probability may thus be overly simplistic and requires further scrutiny. While some studies have found support for Peters’ rule at the level of functional synaptic connectivity for interneurons (Fino & Yuste 2011, Packer et al 2013, Rieubland et al 2014), its general applicability has been refuted, at least for local networks, by dense anatomical reconstructions of retinal (Briggman et al 2011, Helmstaedter et al 2013, Kim et al 2014, Krishnaswamy et al 2015) and cortical circuits (Kasthuri et al 2015, Lee et al 2016). To investigate how this principle applies to long-range projections, a technique that has become widely adopted is subcellular channelrhodopsin-assisted circuit mapping (sCRACM). Here optogenetics is combined with spatially targeted optical stimulation to map the distribution of synaptic currents for a given input (Petreanu et al 2009). While this has been used to show that different presynaptic populations target dendritic subdomains with high specificity (Anastasiades et al 2021, Collins et al 2018, Hooks et al 2013, Yamawaki et al 2019), the extent to which this can be explained and predicted by the distribution of axons and dendrites remains largely an open question.

Whether axons target specific dendritic domains is a particularly important question in the case of layer 5 pyramidal neurons (L5PN), given their central role in several theories of cortical computation (Aru et al 2020, Guerguiev et al 2017, Larkum 2013, Richards et al 2019). For example, the interaction between FF and FB information streams across cortical layers (Larkum et al 2018) is thought to underlie sensory perception (Larkum 2013, Takahashi et al 2020, Takahashi et al 2016) and implement global inference algorithms such as predictive coding (Shipp 2016). These theories all rely heavily on the assumption that FF connections target primarily basal dendrites while FB connections preferentially synapse onto the apical tuft, and would need to be revised should this not be true. This assumption in turn rests on Peter’s rule, but the evidence for this remains circumstantial and a direct examination of Peters’ rule across multiple input pathways to individual neurons remains to be done.

Here we present a comprehensive description of the functional input connectivity to thick-tufted layer 5 (ttL5) pyramidal neurons in medial secondary visual cortex. In particular, we set out to answer three questions: determine the source of input connectivity to ttL5 neurons using monosynaptically restricted rabies tracing (Kim et al 2015, Reardon et al 2016), create a census of subcellular input maps using sCRACM, and test Peters’ rule directly by comparing synaptic input maps to the respective axonal projection maps.

## Results

To ensure recording from a homogeneous neuronal population, we used the Colgalt2-Cre mouse line which specifically labels subcortically projecting, thick-tufted layer 5 (ttL5) neurons (Groh et al 2010, Kim et al 2015). We focused our study on the medial secondary visual cortex (V2M) as higher order cortical regions are likely to receive a broader diversity of long-range inputs than primary sensory cortices. V2M is defined in the Mouse Brain In Stereotaxic Coordinates atlas (Franklin & Paxinos 2007) which can be used to guide viral injections. Furthermore, as this atlas is based on cytoarchitecture, thus V2M can be visually distinguished and selectively targeted in slice recordings, as has previously been done (Galloni et al 2020, Young et al 2021). For whole-brain rabies tracing, on the other hand, we adopted the more recently developed Allen Common Coordinate Framework (CCFv3, (Wang et al 2020a). This atlas allowed us to localize individual neurons within 3D volumes of brain tissue, which is not possible using the Franklin & Paxinos atlas. Within the CCFv3, area V2M corresponds to VISpm, VISam, and RSPagl (Lyamzin & Benucci 2019), all of which are known to be visually responsive (Garrett et al 2014, Powell et al 2020). Treating V2M as a single area for this study was also supported by the observation that axonal projections to VISpm, VISam, and RSPagl are not substantially different (Figure S1, see Methods for details).

### Brain-wide input map to V2M ttL5 pyramidal neurons

We employed a monosynaptically restricted rabies virus approach (Reardon et al 2016, Wickersham et al 2007) to generate a presynaptic input map of V2M ttL5 neurons. Briefly, a mix of adeno-associated viruses carrying floxed N2c G-protein, or TVA-receptor and EGFP genes were injected into V2M of Colgalt2-Cre mice under stereotaxic guidance. Five to seven days later, mCherry expressing EnvA-CVS-N2c-ΔG rabies virus was injected at the same location. After a further 10-12 days, brains were fixed and imaged using serial section 2-photon tomography (Figure S2A). The resultant datasets were registered to the Allen CCFv3 atlas and presynaptic cell bodies were detected and counted using an automated pipeline (see Methods for details).

Cell density maps for an example experiment are shown in Figure 1. Starter cells were scattered across V2M (Figure 1A) while presynaptic input neurons were detected in a broad range of cortical and subcortical areas (Figure 1B,C). We have grouped the most prominent input areas into proximal cortex, distal cortex, and thalamus (Figure 1D). The majority of input cells were found locally in V2M and in the proximal cortical areas VISp and the granular retrosplenial cortex (RSPg, consisting of RSPd and RSPv). Orbitofrontal cortex (ORB) and the anterior cingulate area (ACA) provided the most numerous distal cortical inputs. Interestingly, while most cortical input cells were detected in the granular and infragranular layers, especially layer 5, input from ORB was almost exclusively from layer 2/3 (Figure S2BC).

**Figure 1.**
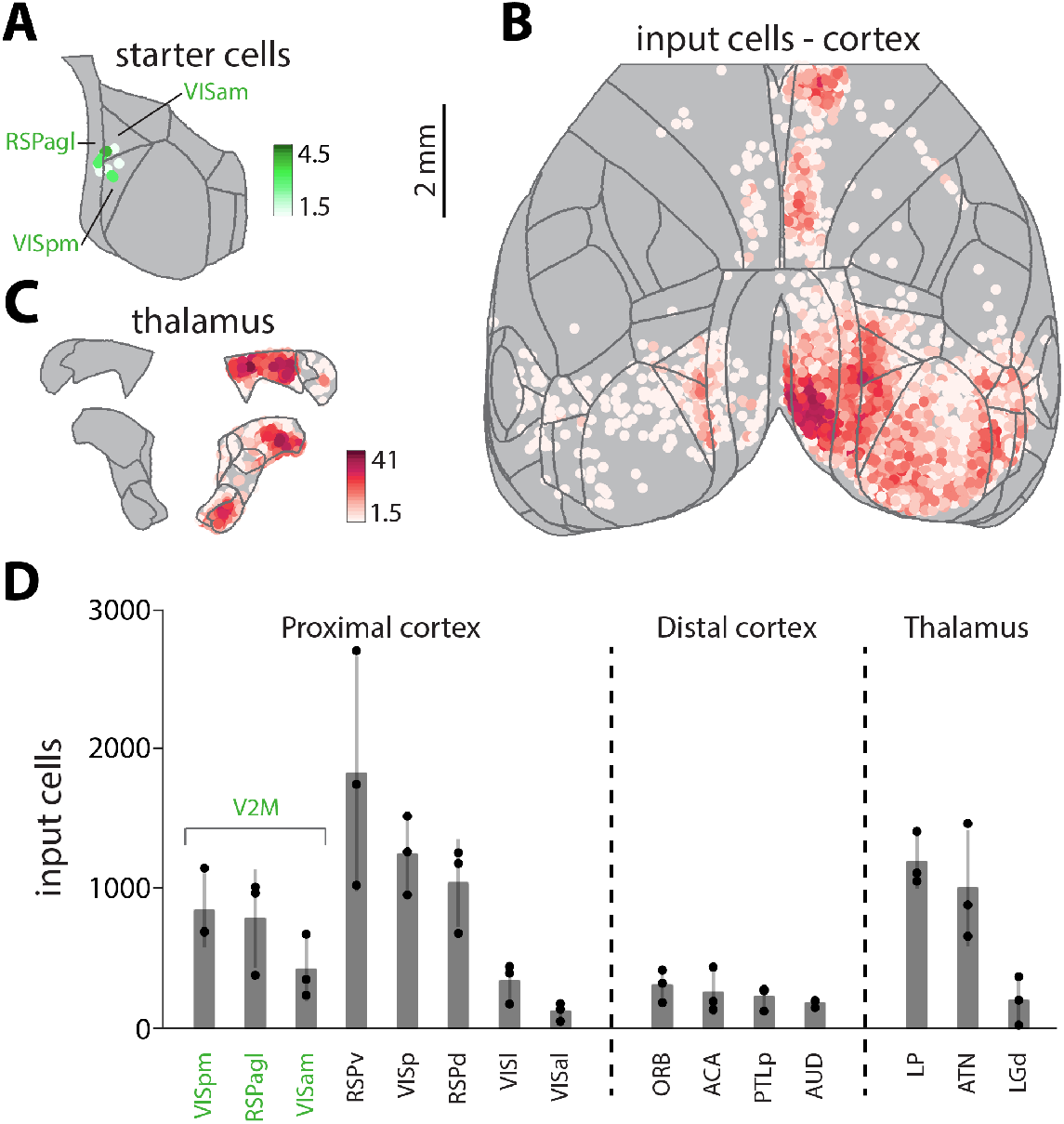
Whole-brain input map to V2M ttL5 neurons. **A.** Starter cell density map from an example experiment. **B.** Cortical input cell density map projected onto the horizontal plane, same experiment as in A. **C.** Thalamic input cell density map projected onto two coronal planes, same experiment as in A. Area names can be found in Figure S3; density scales are in cells / 0.01 mm^2^. **D.** Input cell numbers for the most prominent input areas. Averages, standard deviation, and individual experiments are show.

Prominent thalamic inputs were also observed, originating mainly in the lateral posterior nucleus (LP) and anterior thalamic nuclei (ATN). Comprehensive cell counts for individual experiments can be found in Supplementary table 1.

To understand the organization of inputs onto ttL5 neurons in V2M, we chose to further examine 7 prominent input areas. VISp and V2M for FF input; RSPg, ACA and ORB for cortical FB input; and LP and ATN for thalamic FB connections. We designate local (V2M) input as FF, as ttL5 neurons are considered the outputs of the cortical column, and have very limited local projections.

### Subcellular optogenetic input mapping reveals diverse targeting of dendritic domains by input areas

To determine the spatial distribution of synaptic inputs to ttL5 neurons in V2M, we performed sCRACM experiments from selected input areas identified by the rabies tracing. Following expression of the optogenetic activator Chronos in different input areas (see methods for injection details), we made voltage-clamp recordings (at −70 mV) from tdTomato labelled (Colgalt2-Cre) ttL5 neurons in V2M using acute brain slices. Optical stimulation with a 463 nm laser was spatially targeted using a digital micromirror device (Figure 2A). Sodium and potassium channels were blocked using TTX (1μm) and 4-AP (100 μm) to ensure that evoked currents were restricted to directly stimulated nerve terminals and to enhance presynaptic release, respectively. The stimulus consisted of 24 × 12 spots of light in a 1000 × 500 μm grid aligned to the axis of the apical dendrite of the recorded neuron and covering the whole depth of cortex. We also quantified the total input from a given connection by recording synaptic currents evoked by full-field stimulation. To facilitate comparison between projections, we used the same laser intensity across all experiments.

**Figure 2.**
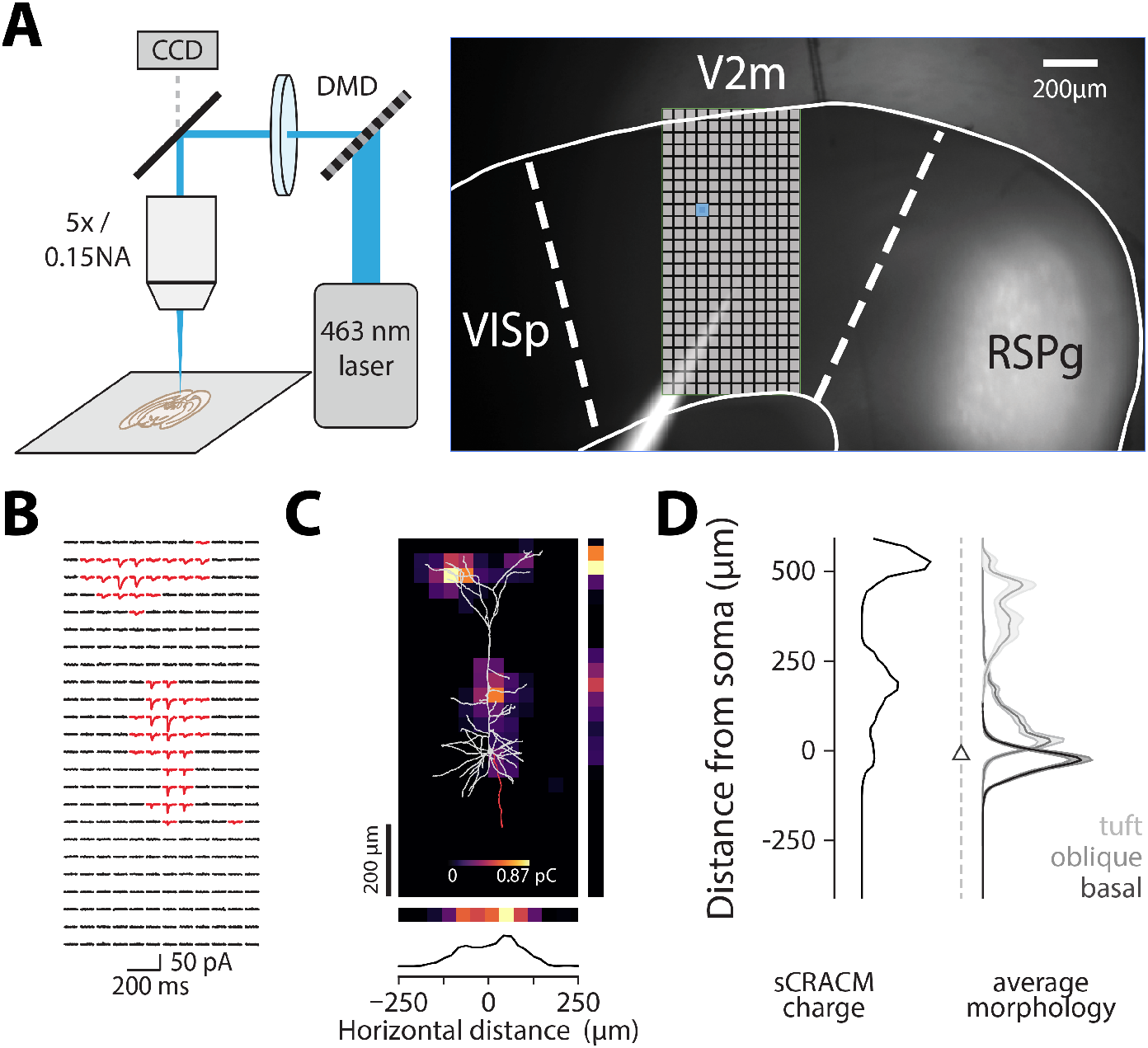
Using sCRACM to map subcellular connectivity. **A.** Experimental setup and micrograph showing a brain slice with Chronos expression in RSPg and recording pipette in V2M. The stimulation grid is overlaid, and an example spot is highlighted in blue. **B**. sCRACM recording of excitatory synaptic currents (red > 7 × baseline S.D.) from an example neuron. **C.** Charge heatmap corresponding to recording in B with the morphology of the recorded neuron overlaid. **D.** Normalized vertical profile of the input map in C. Right: average morphology profile used for dendritic domain deconvolution.

Synaptic strength at each location was estimated by measuring the area of evoked synaptic currents (corresponding to charge; Figure 2B) and creating normalized 2D maps of the spatial distribution of inputs (Figure 2C). Individual maps were then aligned (either to the pia or soma) and averaged. To quantify the spatial location of inputs, we projected the average 2D maps in directions parallel (Figure 2C) or perpendicular to the apical dendrite (Figure 2D). Furthermore, we defined the spatial distribution of the three main dendritic compartments based on 11 morphologically reconstructed Colgalt2-Cre neurons (Figure S4). Basal dendrites were defined as those originating at the soma, oblique dendrites as those originating from the apical trunk before the main bifurcation (including the apical trunk itself), and apical tuft dendrites as those originating after the bifurcation. As all three dendritic compartments have similar spine densities (Romand et al 2011), the horizontal projection of the average morphology was used to separate the contribution of each dendritic domain to the total synaptic input (Figure 2D).

One potential concern when recording distal synaptic currents from a somatic electrode is the effect of attenuation on detectability of currents. In neurons with weaker overall input, this might result in distal currents becoming too small to detect, thus biasing the input map towards the soma. We tested this by examining the correlation between the location of synaptic input and the total synaptic charge evoked by full-field stimulation (Figure S5). No correlation was found for any of the recorded areas, suggesting no detection bias for distal inputs.

#### Primary visual cortex

We first recorded optically evoked synaptic responses arising from VISp axons (n = 9 cells from 6 animals, average soma depth 507 ± 22 μm; Figure 3A). The apical tuft received 42% of the input, with a peak input located 188 μm from the pia (Supplementary table 2). The remaining input was spread between the oblique compartment, receiving 33%, and basal dendrites, receiving 26% of the total input. More of the recorded neurons had the peak input in the apical compartment (n = 5 / 9) while 4 cells lacked apical input (Figure S6). The horizontal input distribution showed a slight medial skew (towards RSPg), most prominent in the oblique (63 μm) and basal compartments (42 μm; Figure S6B). The total synaptic charge measured via the somatic recording following full-field stimulation was 0.93 ± 0.11 pC (Figure 3A). VISp thus provides moderate direct input to ttL5 neurons in V2M, primarily targeting the proximal part of the apical tuft (0.39 pC) with smaller input arriving to the oblique (0.30 pC) and basal (0.24 pC) compartments.

**Figure 3.**
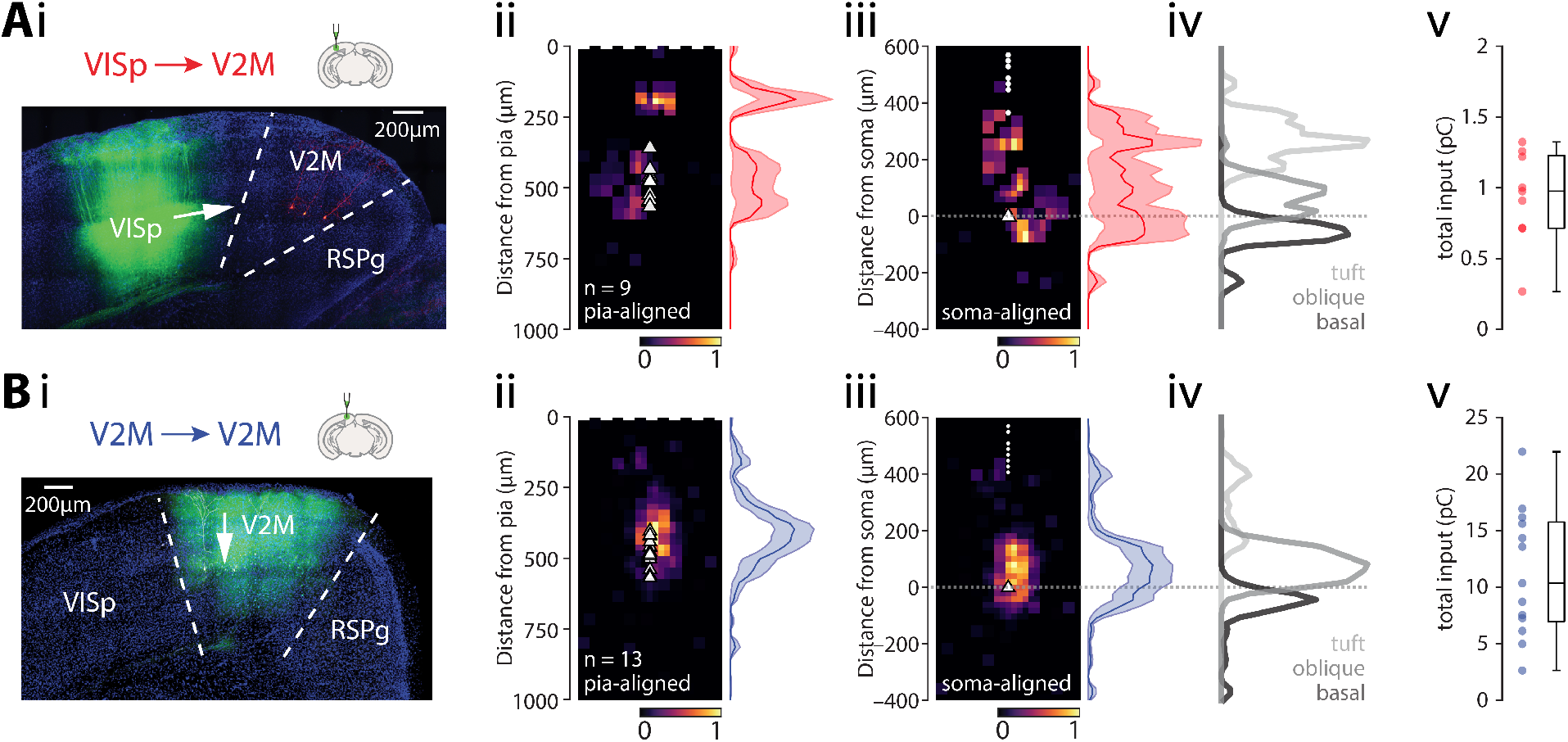
Subcellular connectivity maps of FF input areas. **A.** *i*: confocal image of a representative brain slice (blue = DAPI) showing the injection site in VISp (green) and recorded neurons in V2M (red). *ii*: pia-aligned average sCRACM heatmap for VISp inputs. Triangles represent soma locations. The vertical profile indicates the normalized average and SEM of the input distributions across all recorded neurons. *iii*: Same as in ii but aligned on the soma location. Dots indicate pia locations. *iv*: Normalized input magnitude deconvolved with the average morphology. Dotted line indicates soma location. *v*: Box plot showing total input charge recorded during full-field stimulation. **B.** Same as in A but for Cre-off Chronos injections into V2M.

#### Local input from V2M

To estimate the distribution of local input we used a Cre-off viral strategy, limiting Chronos expression to non-Colgalt2-Cre neurons (n = 13 cells from 4 animals, average soma depth 498 ± 15 μm; Figure 3B). When testing this strategy using the much denser Rbp4-Cre line, we found that only a very small proportion of Cre-positive cells expressed Chronos (3%, Figure S7). The peak input was located close to the soma, at 396 μm from the pia. The oblique compartment received the majority (62%) of this input, with the basal dendrites and apical tuft receiving 24% and 14%, respectively, of the total input (Supplementary table 2). For the majority of recorded neurons, the peak input occurred perisomatically (n = 12 / 13; Figure S6C). The horizontal input distribution showed slight medial bias (−21 μm for all peaks; Figure S6D). The total synaptic charge triggered by full-field stimulation was 11.24 ± 1.56 pC (Figure 3B). Local neurons thus provide large direct input to ttL5 neurons in V2M, primarily targeting the oblique (6.96 pC) compartment, with smaller input arriving to the basal (2.68 pC) and tuft (1.6 pC) compartments.

#### Granular retrosplenial area

Next, we recorded optically evoked synaptic responses arising from RSPg axons (n = 20 cells from 9 animals, average soma depth 503 ± 15 μm; Figure 4A). The overall input displayed a bimodal distribution peaking at 125 and 500 μm from the pia. The apical tuft received 30% of the input, with the oblique compartment receiving 40% and basal dendrites 30% of the total input (Supplementary table 2). For the majority of recorded neurons, the peak input targeted the perisomatic dendrites (n = 18 / 20; Figure S8A). The horizontal input distribution showed slight medial bias (Figure S8B). Total synaptic charge triggered by full-field stimulation was 3.40 ± 0.51 pC (Figure 4A). RSPg thus provides a relatively moderate direct input to ttL5 neurons in V2M, targeting the oblique (1.36 pC), basal (1.04 pC) and apical tuft (1.01 pC) compartments to similar extent.

**Figure 4.**
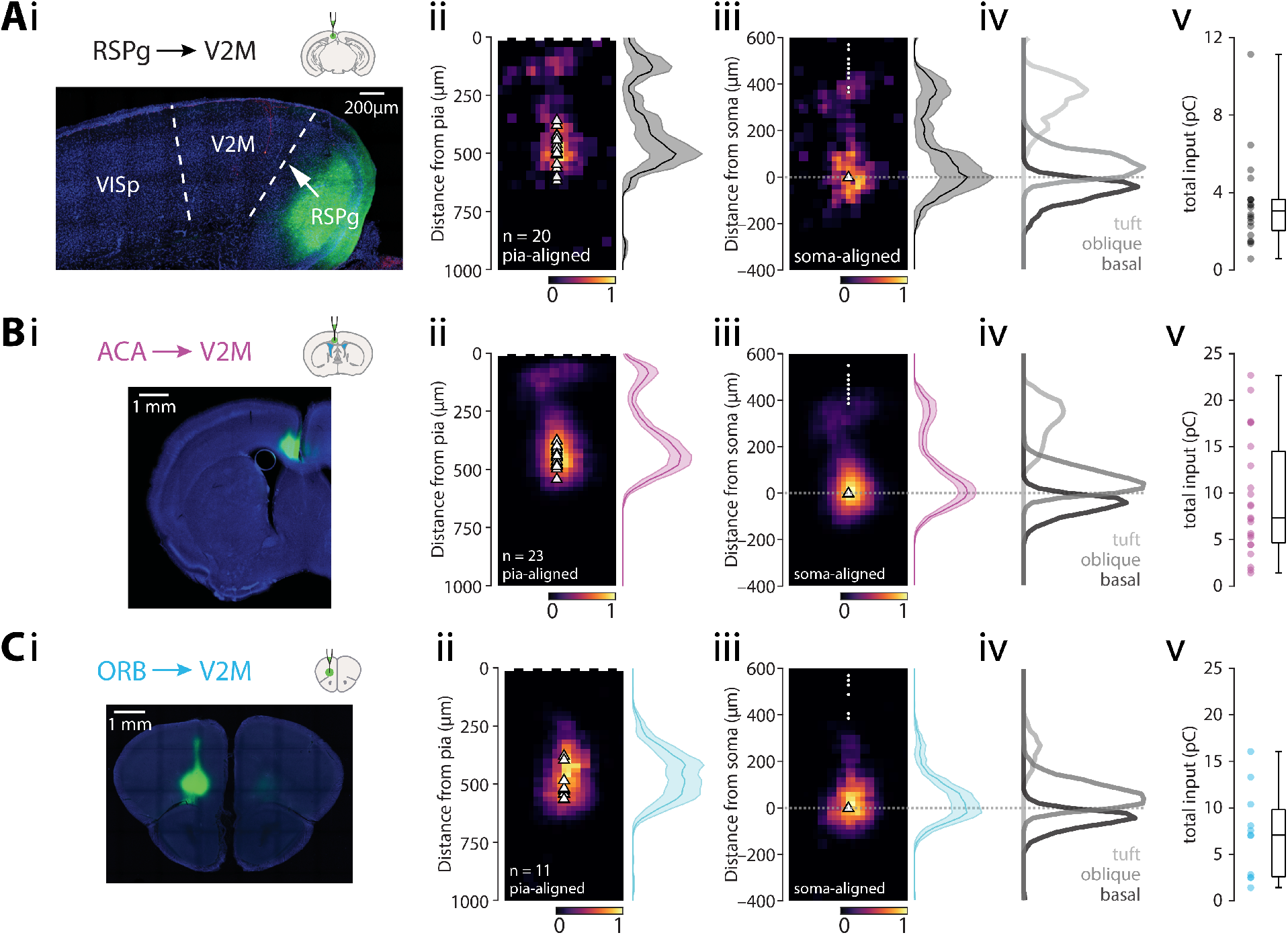
Subcellular connectivity maps of cortical FB areas. **A.** *i:* confocal image of a representative brain slice (blue = DAPI) showing the injection site in RSPg (green). *ii:* pia-aligned average sCRACM heatmap for RSPg inputs. Triangles represent soma locations. The vertical profile indicates the normalized average and SEM of the input distributions across all recorded neurons. *iii:* Same as in *ii* but aligned on the soma location. Dots indicate pia locations. *iv:* Normalized input magnitude deconvolved with the average morphology. Dotted line indicates soma location. *v:* Box plot showing total input charge recorded during full-field stimulation. **B.** Same as in A but for Chronos injections into ACA. **C.** Same as in A but for Chronos injections into ORB.

#### Anterior cingulate area

Next, we recorded optically evoked synaptic responses arising from ACA axons (n = 23 cells from 5 animals, average soma depth 464 ± 9 μm; Figure 4B). The overall input was bimodal, peaking at 83 μm and 438 μm from the pia. The apical tuft received 25% of the input, with the oblique compartment receiving 45% and basal dendrites 30% of the total input (Supplementary table 2). The majority of recorded neurons had the peak input located perisomatically (n = 22 / 23; Figure S8C). The horizontal input distribution showed no medio-lateral bias (Figure S8D). The total synaptic charge triggered by full-field stimulation was 9.46 ± 1.32 pC (Figure 4B). ACA thus provides a large direct input to ttL5 neurons in V2M, primarily targeting the oblique (4.22 pC) compartment with smaller input arriving to the basal (2.83 pC) and most distal part of the apical tuft (2.41 pC).

#### Orbitofrontal cortex

Optically evoked synaptic responses arising from ORB axons (n = 11 cells from 3 animals, average soma depth 521 ± 19 μm; Figure 4C) showed a strong perisomatic bias, with a peak at 417 μm from the pia. The apical tuft received only 9% of all input, with the oblique compartment receiving 57% and basal dendrites 35% of the total input (Supplementary table 2). This distribution was also highly homogeneous across neurons, with almost all recorded neurons having their peak input in the perisomatic region (n = 11/11; Figure S8E). The horizontal input distribution showed no lateral bias (Figure S8F). The total synaptic charge triggered by full-field stimulation was 7.16 ± 1.43 pC (Figure 4C). ORB thus provides a large direct input to ttL5 neurons in V2M, primarily targeting the oblique (4.05 pC) and basal (2.47 pC) compartments, with slight input arriving to the proximal part of the apical tuft (0.63 pC).

#### Anterior thalamic nuclei

Next, we recorded optically evoked synaptic responses from thalamic axons, starting with the ATN (n = 8 cells from 3 animals, average soma depth 435 ± 15 μm; Figure 5A). This input had peaks at both 104 μm and 333 μm from the pia, with the apical tuft receiving the majority (75%) of the input, while the oblique compartment received 17% and basal dendrites a mere 8% of the total input (Supplementary table 2). The majority of recorded neurons had the peak input in the tuft compartment (n = 6 / 8) and while all cells had some tuft input, in 2/8 cells the input peak was located perisomatically (Figure S9A). The horizontal input distribution showed a medial bias (Figure S9B). The total synaptic charge triggered by full-field stimulation was 2.48 ± 0.54 pC (Figure 5A). ATN thus provides a moderate direct input to ttL5 neurons in V2M, primarily targeting the more distal part of the apical tuft (1.86 pC) while the oblique (0.42 pC) and the basal (0.21 pC) compartment received less input.

**Figure 5.**
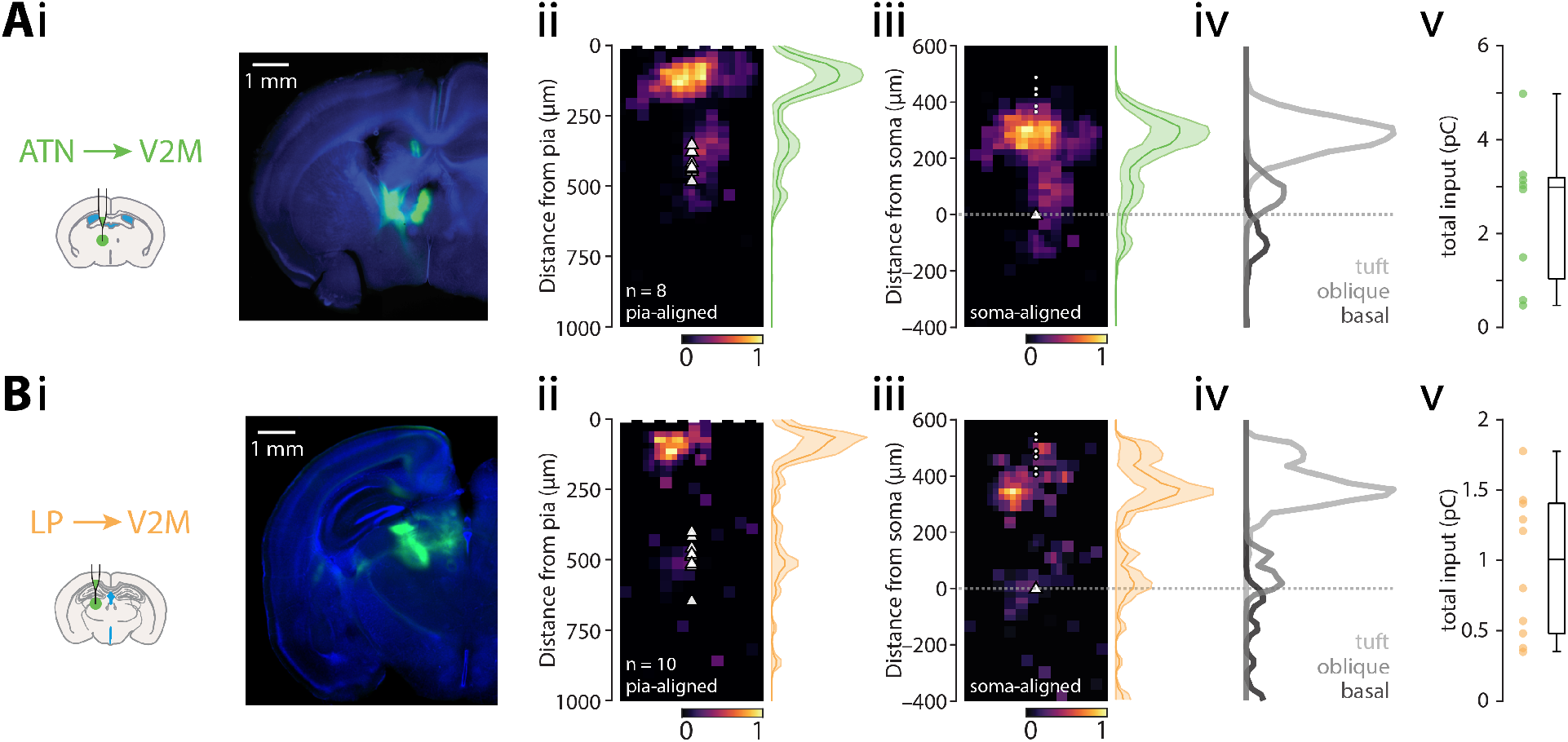
Subcellular connectivity maps of thalamic input areas. **A.** *i:* confocal image of a representative brain slice (blue = DAPI) showing the injection site in ATN (green). *ii:* pia-aligned average sCRACM heatmap for ATN inputs. Triangles represent soma locations. The vertical profile indicates the normalized average and SEM of the input distributions across all recorded neurons. *iii:* Same as in *ii* but aligned on the soma location. Dots indicate pia locations. *iv:* Normalized input magnitude deconvolved with the average morphology. Dotted line indicates soma location. *v:* Box plot showing total input charge recorded during full-field stimulation. **B.** Same as in A but for Chronos injections in LP.

#### Lateral posterior nucleus of the thalamus

Lastly, we recorded optically evoked synaptic responses arising from LP axons (n = 10 cells from 4 animals, average soma depth 500 ± 23 μm; Figure 5B). Due to excessive retrograde labelling resulting in direct photocurrent in the recorded V2M cells, a 1:10 dilution of virus was used for these injections and the absolute value of the evoked input is thus likely an underestimate. As with ATN axons, the LP input was strongly biased towards the most superficial part of the cortex and peaked at 63 μm from the pia. The apical tuft received the vast majority (75%) of the input, with the oblique compartment receiving 15% and basal dendrites 10 % of the total input (Supplementary table 2). Most recorded neurons had the peak input in the tuft compartment (n = 9/10; Figure S9C). The horizontal input distribution showed lateral bias (Figure S9D). The total synaptic charge triggered by full-field stimulation was 0.97 ± 0.16 pC (Figure 5B). LP thus provides modest direct input to ttL5 neurons in V2M, primarily targeting the most distal part of the apical tuft (0.72 pC) compartment with smaller input arriving to the oblique (0.14 pC) and the basal (0.10 pC) compartments.

### Comparison of anatomical and functional connectivity maps

Having determined the spatial distribution of synapses for the main input areas, we next sought to directly compare this to what would be predicted from axo-dendritic overlap (i.e. from Peters’ rule). To determine axonal projection patterns from input areas to V2M, we have imaged the Chronos-eGFP labelled axons in a subset of the brain slices used for the sCRACM experiments using confocal microscopy.

The spatial distribution of axons followed three basic patterns. Axons from VISp and ORB were densest in layer (L) 2/3 and L5 while little projection was apparent in L1, reminiscent of the classical FF projection pattern (Figure 6A). In contrast, axons from RSPg and ACA showed an FB-like pattern with dense labelling in the middle part of L1 followed by sparse labelling in L2 and diffuse axons in layers 3, 5 and 6 (Figure 6B). The final group, which consists of the thalamic projections from LP and ATN, showed the classical FB pattern strongly innervating the external part of L1, with a secondary peak in L3, but little or no projections in layers 2, 5 and 6 (Figure 6C).

**Figure 6.**
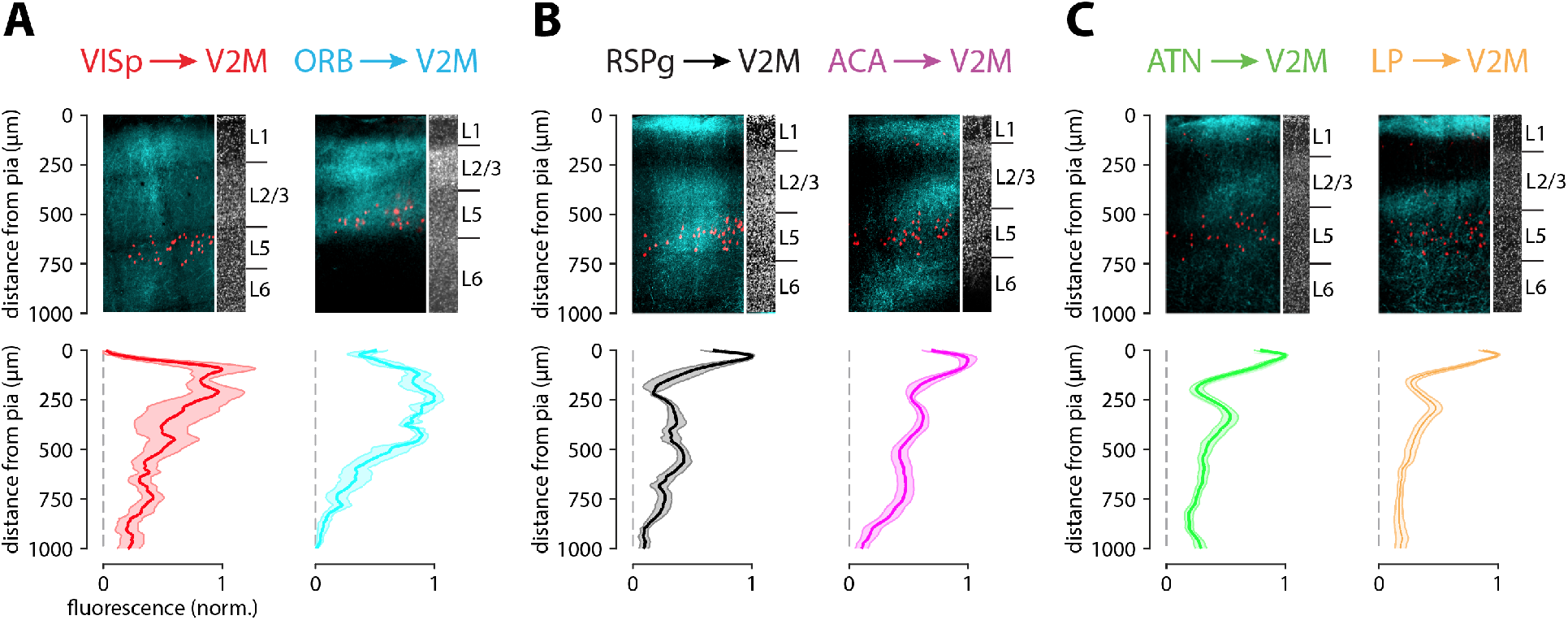
Axonal projection densities from different input areas. **A-C.** Top: example confocal images from V2M showing axonal projections from six input areas (cyan) and Colgalt2-Cre cell bodies (red). The corresponding DAPI staining shows variation in laminar depth. Bottom: average projection density profiles across the cortical depth averaged across 3-5 injections.

To accurately estimate morphological overlap between axons and dendrites, we multiplied the axonal projection maps with the average dendritic morphology, resulting in the predicted input distribution one would expect to see based on Peters’ rule. When overlaying this with the pia-aligned vertical sCRACM maps, the alignments between functional synapses and the axo-dendritic maps were diverse (Figure 7A). For some regions, like ORB perisomatic and LP tuft inputs, a clear correspondence could be seen between predicted and measured input distributions. A lesser degree of overlap can be seen in the VISp perisomatic or ACA tuft inputs. For other inputs, however, strong functional input could be detected where there is little overlap between dendrites and axons, such as at VISp tuft inputs. This stood in stark contrast to the ORB projection, for which the opposite was true, and apical regions of dense morphological overlap of axons and dendrites resulted in no functional input.

**Figure 7.**
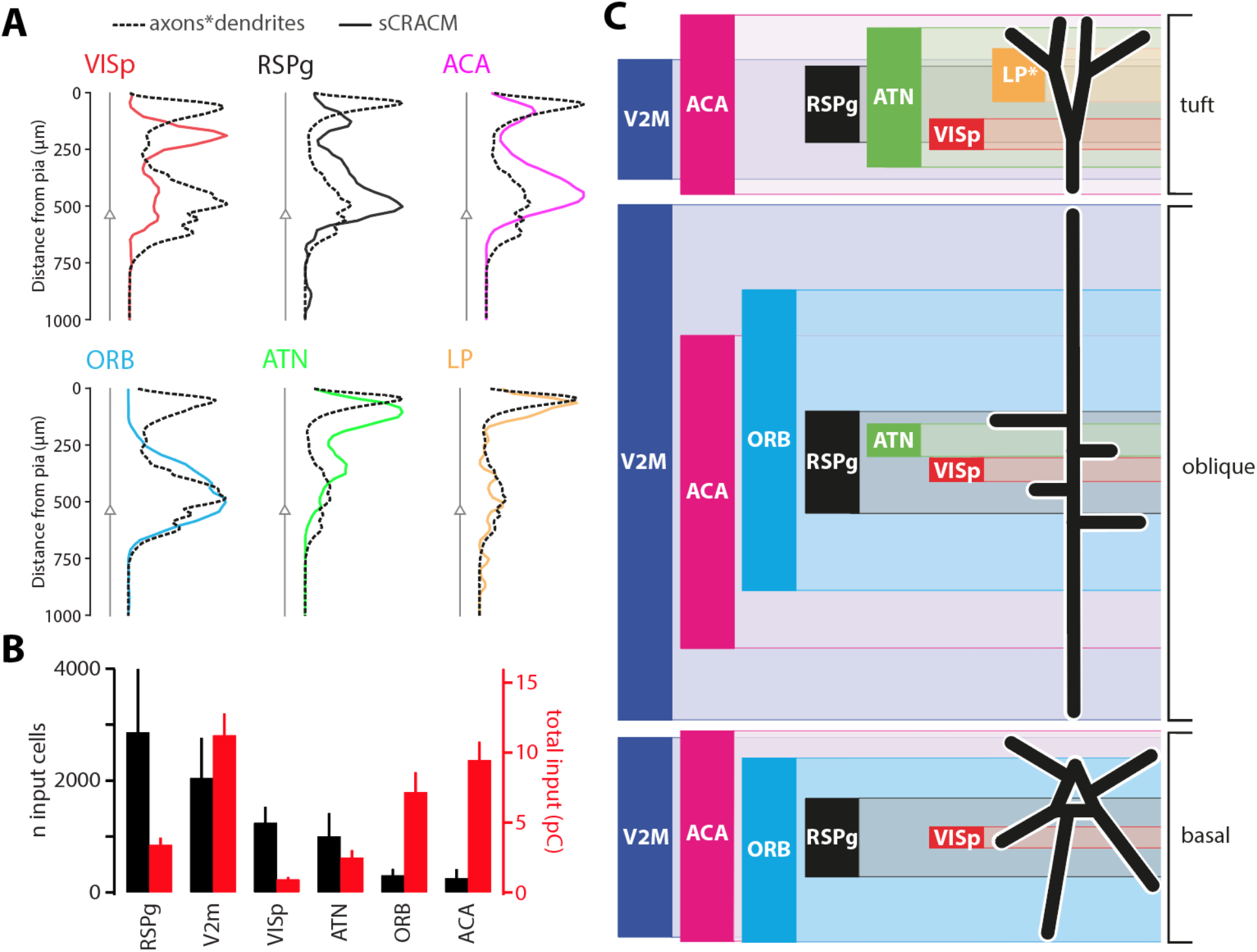
Comparison of different input maps. **A.** Axonal density distributions multiplied with dendritic morphology (dotted black lines) overlaid with pia-aligned synaptic input distributions (coloured lines). Six input areas are shown. Triangles represent soma location in average morphology. **B.** Number of input cells across 6 areas established by rabies tracing (black) and total input charge recorded during full-field optogenetic stimulation (red). **C.** Schematic of excitatory synaptic input map to ttL5 pyramidal neurons. The height of bars represents input strength, while the centre of each bar is aligned to the peak of the sCRACM map. The tuft input map was generated from pia aligned maps while oblique and basal maps from soma aligned maps. NB: LP input magnitude (*) is likely underestimated due to the lower virus titer used.

Next we examined the correspondence between the anatomical input connectivity obtained from rabies tracing and the functional connectivity measured by total synaptic input. The number of rabies-labelled input neurons showed a strong contribution from RSPg and V2M, and modest input cell numbers for the more distal cortical regions (e.g. ACA, ORB). The total synaptic input, however, shows no correlation with these numbers (p = 0.8, r = −0.14, Spearman correlation, Figure 7B), with modest synaptic input from RSPg and most input arriving from V2M, ACA and ORB. Taken together, these results show clear specificity of dendritic targeting by brain-wide connections, with only a loose adherence to Peters’ rule for most inputs as well as large differences between anatomical and functional connectivity measured by rabies tracing and optogenetic stimulation, respectively.

## Discussion

Using an array of techniques for long-range circuit dissection, we have comprehensively mapped the location and dendritic targets of inputs to ttL5 neurons in the medial secondary visual cortex in mice. This allowed us to determine the dendritic targets of FF and FB connections and to make a direct assessment of Peters’ rule for brain-wide connections.

The whole-brain input map generated via rabies tracing was qualitatively similar to previous results from the primary visual cortex (Kim et al 2015). Axonal projections from the rabies-identified input regions broadly followed the expected pattern, with FB projections being biased toward L1 and FF areas toward the deeper layers (D’Souza et al 2020, Harris et al 2019, Rockland & Pandya 1979). Accordingly, L1 was densely innervated by higher-order cortical areas like RSPg and ACA as well as the secondary thalamic nuclei (LP, ATN). Interestingly, ORB, a higher-order cortical region, displayed a projection pattern associated with FF areas. Additionally, the majority of ORB projection neurons were found in L2/3, another feature of FF connectivity. ORB thus seems to be an exception in terms of FB axonal projections.

Compared to the axonal projection patterns, synaptic input maps showed a remarkable degree of heterogeneity. Morphological averaging was necessary as the sCRACM recordings in our dataset did not have paired reconstructions for every cell. While it is possible that comparing the individual axon, dendrite, and synaptic profiles on a single-cell basis would have given slightly more accurate results, the overall pattern of functional input from each input region was mostly consistent across cells. Morphologies of ttL5 neurons are likewise highly stereotypical. Furthermore, the axon projection patterns used for evaluating Peters’ rule were measured from a subset of the same slices used for the sCRACM recordings, further supporting the direct comparison of the predicted input maps with those recorded functionally. The discrepancy resulting from averaging is thus likely to be low. Indeed, any smoothing resulting from averaging of morphologies, axonal projections or sCRACM maps would only increase overlap, and thus bias the results in favor of adhering to Peters’ rule. In contrast to this, several connections showed only weak correspondence between predicted and observed input maps. The connection from VISp, which is FF by definition and as such is assumed to primarily target perisomatic dendrites (Larkum 2013), was instead biased towards the apical tuft. Conversely, while the axons from ORB had a FF-like projection pattern, they synapsed almost exclusively with basal and oblique dendrites. Other recorded areas (e.g., RSPg, ACA, ATN) showed some degree of conformity to Peters’ rule, yet still with significant differences in the proportion of synapses generated in the different regions of high axonal projections. The only area where the correspondence was remarkable and Peter’s rule held fully was LP.

Comparing anatomical connectivity obtained by rabies tracing to functional connectivity obtained by full-field optogenetic stimulation of axons revealed large and unexpected differences. When evaluating this finding, it is important to consider a few technical caveats which might bias this comparison. First, while we have not found false-positive areas (i.e., every area revealed by rabies tracing and tested by sCRACM provided input), there is considerable debate regarding the quantitative accuracy of rabies tracing (Rogers & Beier 2021). Second, the magnitude of optogenetically evoked input depends on the number of presynaptic cells covered by viral injection. To facilitate comparison with the rabies labelling, we aimed to maximise coverage of each area by making several injections targeted to the locations with the highest density in the rabies data. It is unlikely that these technical caveats could alone account for the remarkable discrepancy between anatomical and functional input magnitudes. There are several other possible explanations for this difference. First, there may be differing convergence of connectivity between input areas. For example, low convergence in inputs with strong rabies labelling, with a unique mapping of selected pre- and postsynaptic cells, could result in relatively weaker sCRACM input (like VISp and RSPg). Meanwhile, strong synaptic currents relative to small rabies-labelled populations (like ORB and ACA) may be explained by higher convergence. Such connections might be less discerning of their targets in order to convey more general contextual or state-specific information. Second, there may be a difference in the strength of individual synapses not reflected in rabies efficiency, with sparsely labelled input areas like ORB having relatively strong synapses, while rabies-dense areas like RSPg may send a large number of weaker synapses. A third contributing factor could be the recently reported activity-dependence of rabies transmission (Beier et al 2017). The apparent sparsity of some input areas (like ORB and ACA) could thus arise from having very low activity. Conversely, to result in extensive rabies labelling, VISp and RSPg should provide high activity input.

We used a novel approach to allocate sCRACM input to specific dendritic compartments by deconvolving the synaptic input maps using morphological ttL5 reconstructions in which we manually labelled tuft, oblique, and basal compartments. Our results provide a complex picture regarding the possible interaction of FF and FB inputs (Figure 7C). Functionally, V2M has been linked to visual motion processing (Sun et al 2009) and is thought to take part in navigation and spatial processing as part of the dorsal stream (Glickfeld & Olsen 2017). Thalamic FB input, which targets almost exclusively the apical tuft, arrives from multiple higher-order nuclei. Parts of ATN receive strong vestibular input (Rancz et al 2015) and, together with RSP form a central part of the head-direction system (Taube 2007, Velez-Fort et al 2018). It is thus likely that spatial and multisensory contextual information carried by LP inputs (Roth et al 2016) interacts with FF visual input in the tuft compartment. However, the role of tuft integration is likely to differ from primary sensory cortices, considering that ttL5 neurons in the secondary visual cortex have substantially different integrative properties (Galloni et al 2020). Contrary to thalamic input, cortical FB inputs target all three dendritic domains. Perhaps surprisingly, the strongest of these are ORB and ACA, which can interact with the local FF input at the level of oblique and basal dendrites. Frontal cortices in general are involved in decision making and executive control of behavior (Hamilton & Brigman 2015), and ORB in particular has been shown to encode spatial goals (Feierstein et al 2006). ACA, meanwhile, can directly regulate visually evoked responses and sensory discrimination in the primary visual cortex (Zhang et al 2014), and contributes to learning to predict sensory input to primary visual cortex (Fiser et al 2016). Their precise roles in the functioning of V2M, however, remains unknown. The strong input received by oblique dendrites is particularly important, as this compartment was shown to strongly affect L5 excitability (Schaefer et al 2003) and can gate information flow from the apical dendrites (Jarsky et al 2005). While FB inputs targeting the apical tuft have been suggested to act as a general gain control for ttL5 neurons, it is thus possible that ORB and ACA perform a similar gating function, but perhaps with more specificity regarding input identity or dendritic branches. Indeed, their particularly strong targeting of oblique dendrites might allow both fine-level control of plasticity for FF synapses at these dendrites, while simultaneously allowing them to exert gating control over input from both the thalamic nuclei and VISp, which strongly project to the tuft.

In general, our results show that while the classification of areas as FF or FB can be based on axonal projections (albeit with exceptions, such as ORB), macroscopic projectomes do not predict cell-type level input location, and individual connections do not follow clear rules associated with their position above or below the target area in a cortical hierarchy. Similarly, while rabies tracing from a population of starter cells is an effective tool to study the general wiring diagram, the proportion of input neurons thus estimated is likely to give a poor estimation of functional input strength. Finally, the location and possible interactions between FF and the broad range of FB inputs as well as their specific information content suggests that ttL5 neurons may be adopting a multitude of integrative strategies that are more complex than those previously suggested.

## Materials and Methods

### Animals

All animal experiments were prospectively approved by the local ethics panel of the Francis Crick Institute (previously National Institute for Medical Research) and the UK Home Office under the Animals (Scientific Procedures) Act 1986 (PPL: 70/8935). All surgery was performed under isoflurane anesthesia, and every effort was made to minimize suffering. Transgenic mice were used: Tg(Colgalt2-Cre)NF107Gsat (RRID:MMRRC_036504-UCD, also known as Glt25d2-Cre) were crossed with the Ai14 reported line expressing tdTomato (RRID:IMSR_JAX:007908). Additionally, Tg(Rbp4-Cre)KL100Gsat/Mmucd (RRID:MMRRC_031125-UCD) mice were used to establish the efficacy of the cre-off approach. As only male mice are transgenic in the Colgalt2-Cre line, all experiments were done on male animals. Animals were housed in individually ventilated cages under a 12 hr light/dark cycle.

### Viruses

EnvA-CVS-N2c^ΔG^-mCherry rabies virus, and adeno associated viruses expressing TVA and EGFP (AAV8-EF1a-flex-GT), N2c glycoprotein (AAV1-Syn-flex-H2B-N2CG), or Cre-OFF Chronos-GFP (AAV1-EF1-CreOff-Chronos-GFP) were a generous gift of Molly Strom and Troy Margrie. Chronos-GFP (also called ShChR) expressing adeno associated virus (rAAV1-Syn-Chronos-GFP) was obtained from UNC Vector Core.

### Surgical procedures

Surgeries were performed on mice aged 3–8 weeks using aseptic technique under isoflurane (2–4%) anesthesia and analgesia (meloxicam 2 mg/kg and buprenorphine 0.1 mg/kg). The animals were head-fixed in a stereotaxic frame and a small hole (0.5–0.7 mm) was drilled in the skull above the injection site. Virus was loaded into a glass microinjection pipette (pulled to a tip diameter of around 20 μm) and pressure injected into the target region at a rate of 0.4 nl/s using a Nanoject III delivery system (Drummond Scientific). To reduce backflow, the pipette was left in the brain for approximately 5 min after completion of each injection.

For rabies virus tracing experiments, a 1:2 mixture of TVA and N2c glycoprotein expressing cre-dependent AAVs (10-20 nL) was injected at stereotaxic brain coordinates (ƛ − 0.8 mm, ML 1.6 mm, DV 0.6 mm). Rabies virus (50-100 nL) was injected 5-7 days later at the same site. Ten to twelve days later, animals were transcardially perfused under terminal anesthesia with cold phosphate-buffer (PB, 0.1 M) followed by 4% paraformaldehyde (PFA) in PB (0.1 M).

For the sCRACM experiments, Chronos-GFP expressing AAV was injected into one of the identified presynaptic regions. The virus was allowed to express for at least 3 weeks before acute brain slice preparation. The range of stereotaxic coordinates for each region are listed in Supplementary table 3. The injected virus was Chronos-GFP for every region except V2M, where Cre-OFF Chronos-GFP was instead used to avoid expression in the recorded Colgalt2-Cre neurons. For some of the injections in LP, the Chronos-GFP virus was diluted by 10-fold in sterile cortex buffer before injection.

### Data acquisition and analysis for rabies tracing experiments

Brain samples were embedded in 4-5% agarose (Sigma-Aldrich: 9012-36-6) in 0.1M PB and imaged using serial two-photon tomography (Han et al 2018, Osten & Margrie 2013, Ragan et al 2012). Eight optical sections were imaged every 5 μm with 1.2 μm × 1.2 μm lateral resolution, after which a 40μm physical section was removed. Excitation was provided by a pulsed femto-second laser at 800 nm wavelength (MaiTai eHP, Spectraphysics). Images were acquired through a 16X, 0.8 NA objective (Nikon MRP07220) in three channels (green, red, blue) using photomultiplier tubes. Image tiles for each channel and optical plane were stitched together with an open-source software written in MATLAB (https://github.com/SainsburyWellcomeCentre/StitchIt). For cell detection, full resolution images were first filtered with a Gaussian blur (sigma = 1) using Fiji (ImageJ 1.52e) to reduce imaging noise. The open-source package “cellfinder” (Tyson et al 2020) was used for cell candidate detection then classification. Automated mouse atlas propagation (Niedworok et al 2016) was used for registration and segmentation on brain samples down-sampled to 10 μm voxels (to match the resolution of the Allen CCFv3; (Wang et al 2020a). Cell coordinates were similarly down-sampled to 10 μm and the number of cells was counted for each segmented area. For cell density visualisation, cell coordinates were reverse-transformed onto the Allen CCFv3 space using the open source registration tool, Elastix (Klein et al 2010) and were projected onto a 2D matrix in 10 μm / pixel resolution.

### Acute slice preparation and electrophysiological recordings

Adult mice were deeply anaesthetised with isoflurane and decapitated. The brain was rapidly removed and placed in oxygenated ice-cold slicing solution containing (in mM): 125 sucrose, 62.5 NaCl, 2.5 KCl, 1.25 NaH_2_PO_4_, 26 NaHCO_3_, 2 MgCl_2_, 1 CaCl_2_, 25 dextrose; osmolarity 340–350 mOsm. The cerebellum and frontal cortex were removed manually with a coronal cut using a single-edged razor blade and the rostral surface was affixed to a metal platform with cyanoacrylate glue. Coronal slices (300 μm thick) between 2.6 and 3.5 mm posterior to bregma were prepared using a vibrating blade microtome (Leica VT1200S). Slices were kept submerged in artificial cerebrospinal fluid (ACSF, containing in mM: 125 NaCl, 2.5 KCl, 1.25 NaH_2_PO_4_, 26 NaHCO_3_, 1 MgCl_2_, 2 CaCl_2_, 25 dextrose; osmolarity 308–312 mOsm) at 35°C for the first 30–60 min after slicing, then at room temperature (22°C). All solutions and chambers were continuously bubbled with carbogen (95% O2 / 5% CO2).

The topology of the cortical mantle in the region of V2m is curved and varies in thickness along the antero-posterior axis. In most coronal sections, the apical dendrites of L5 neurons were thus at a slight angle relative to the slicing surface. This angle was minimized by slicing the brain with a slight backward angle relative to the coronal plane. Additionally, to avoid recording from neurons with cut apical dendrites, slices were placed such that the apical dendrites could be seen to descend at a shallow angle into the slice. Where possible, the fluorescently-labelled apical dendrites were also visually inspected along their full path from soma to pia. Furthermore, many neurons were successfully filled with biocytin during recording, making it possible to verify the integrity of the apical dendrite after the recordings. The observed neurons were all found to have an intact apical trunk with tuft dendrites extending to the pia. However, we can’t exclude that a small fraction of finer dendrites (including both basal and apical tuft) extending towards the slice surface may have been partially cut in the process.

For recordings, individual slices were perfused in the recording chamber at a rate of approximately 6 mL/min with ACSF at room temperature (22°C), continuously bubbled with carbogen. To prevent axonal spike propagation and enhance responses to optical stimulation, 1 μM tetrodotoxin (TTX) and 100 μM 4-aminopyridin (4-AP) were added to the recording ACSF. This ensured that any light-evoked responses were direct monosynaptic responses resulting from stimulation of Chronos-expressing axon terminals, rather than from passing axons terminating in unknown locations on the dendrites.

Filamented borosilicate thick-walled glass micropipettes were pulled and heat-polished using a two-stage horizontal puller (Zeitz DMZ Universal Electrode Puller) to obtain an electrode resistance of 3–6 MΩ. The glass electrodes were filled with internal solution optimized for voltage clamp recordings, containing (in mM): 120 CsMeSO_3_ (CH_3_O_3_SCs), 3 CsCl, 10 HEPES, 1 EGTA, 4 Na_2_ATP, 0.3 NaGTP, 5 Na_2_-phosphoreatine (C_4_H_8_N_3_O_5_PNa_2_), 3.5 QX-314 chloride, 0.5 % (w/v) biocytin hydrochloride, 50 μM Alexa Fluor 488 hydrazide; osmolarity 290–295 mOsm; pH adjusted to 7.3 with CsOH.

Visually guided whole-cell patch-clamp recordings from tdTomato-labelled Colgalt2-Cre neurons in V2M were performed using a Scientifica SliceScope Pro 3000 microscope equipped with a 40x/0.8 NA objective and an infrared (IR) Dodt Gradient Contrast system. The epifluorescence system used to visualize fluorescent neurons was a CoolLED pE-4000 illumination system with a 550 nm peak excitation wavelength. To avoid stimulating Chronos-expressing axons, epifluorescent illumination was kept to a minimum during selection of cells to record. Recordings were made with a Multiclamp 700B amplifier (Molecular Devices) in voltage-clamp configuration with a holding potential of −70 mV. Filtered signals (8kHz low-pass) were digitized at 20 kHz with a National Instruments DAQ board (PCIe-6323). Acquisition and stimulus protocols were generated in Igor Pro (Wavemetrics) with the NeuroMatic software package (Rothman & Silver 2018). Throughout each recording, series resistance compensation was applied and set to the highest value possible without inducing oscillations in the cell (typically between 40 and 75%). Recordings wtih series resistance larger than 40 MΩ were excluded.

### Patterned optogenetic stimulation

Optical stimulation was implemented using a digital micromirror device (DMD) with a 463 nm laser (laser-coupled Polygon 400, Mightex Systems). The stimulus consisted of a 1000 × 500 μm grid divided into 24 × 12 spots of light (41.7 μm × 41.7 μm square) delivered through a 5x/0.15 NA dry objective (Olympus MPlanFL N). The grid was approximately centered on the soma being recorded from, aligned to the pia orthogonal to the apical dendrite. For each individual spot, the laser power was measured at the specimen plane using a PM100D (Thorlabs) optical power meter equipped with a S121C sensor (Thorlabs). The laser output associated with each spot was adjusted to obtain a measured power of approximately 300 μW (173 mW/mm^2^).

Optical stimuli were delivered for 1 ms at 0.1 Hz in a pseudo-random sequence designed to maximise the distance between consecutive spots and the time between stimulation of neighbouring spots. Each recording trial consisted of a single repetition of all 288 stimuli followed by a full-field stimulus, in which all stimulation spots were illuminated simultaneously for 1 ms. For each cell, 5-20 trials were recorded, with 30s pauses between trials, making the interval between consecutive stimulation of the same spot approximately 60s. Following each recording, an image was taken to record the location of the recorded cell (filled with Alexa Fluor 488) relative to the stimulation grid. This was used during analysis to align the recorded sCRACM heatmap with the location of the pia or soma.

### Immunohistochemistry & morphological reconstructions

After recording, slices were fixed overnight at 4°C in a 4% paraformaldehyde solution and were subsequently kept in PBS. Slices were stained with DAPI (5 μg/mL) for 10 min, mounted on glass slides and images were acquired with either a confocal microscope for high-resolution images (Leica SP5; objective: 20x/0.7NA or 10x/0.4NA; pinhole size: 1 airy unit) or a slide scanner for visualizing injection sites (Olympus VS120, objective: 4x/0.16NA). Image processing was done with the FIJI software package (Schindelin et al 2012). For the detailed morphological analysis, a subset of neurons, selected based on the quality and completeness of staining, was reconstructed in full through the LMtrace service of https://ariadne.ai/lmtrace.

### Comparison of axonal projection patterns to VISam, VISpm and RSPagl

We have obtained layer-wise axonal projection data from the 6 input areas (ACA n=33, ATN n=11, LP n=10, ORB n=11, RSPg n=17, VISp n=60) from the Allen Mouse Brain Connectivity database (© 2011 Allen Institute for Brain Science. Allen Mouse Brain Connectivity Atlas. Available from: https://connectivity.brain-map.org/). Projection energy distributions were qualitatively similar across target areas (Figure S1). Quantitatively, the data followed the same layer-wise pattern across the three target areas (p > 0.05, 2-way ANOVA with Tukey’s post hoc test) with the exception of ACA (RSPagl vs VISpm p = 0.001; VISam vs VISpm p = 0.016). When only wild-type data was considered, no statistical difference between target areas was detected (p > 0.05, 2-way ANOVA with Tukey’s post hoc test; n = ACA 5, ATN 4, LP 2, ORB 2, RSPg 2, VISp 21).

### Data analysis

Analysis and data visualization were performed with custom macros and scripts written in Igor Pro and MATLAB (Mathworks). Unless otherwise specified, all reported data values refer to the mean ± standard error (SEM). Recordings were not corrected for liquid junction potential.

Recordings were baselined in a 40 ms window before each stimulus and averaged across trials, and the peak and area of the evoked currents were measured in a 50 ms window after the stimulus. Stimulus spots for which the peak current was lower than seven times the standard deviation of the baseline noise were scored as zero.

For some of the injection sites, a degree of retrograde transport of the Chronos virus was noted in areas outside of the primary injection site, including in V2M. A few recorded Colgalt2-Cre neurons in V2M were thus found to be intrinsically expressing Chronos. This was easily detected in the recordings by an instantaneous inward current at the onset of laser stimulation, in contrast to the 4-5 ms delay between stimulus onset and sCRACM current observed normally. Any cells with no such delay were excluded from the analysis.

Because peak response amplitudes varied between cells and preparations, to obtain average input distributions from a presynaptic population, the heatmap for each cell was normalised to the peak EPSC value for that cell. Heatmaps were then aligned horizontally by soma location and then vertically by either soma or pia location before averaging each pixel across cells. For each cell, the soma could be localised to within one quadrant of a given stimulation spot. When averaging across cells, the effective sampling resolution (i.e. the pixel dimension) for the average sCRACM heatmaps and related horizontal and vertical projections were thus approximately 20.8 μm (equal to half of the stimulus spot size, i.e. 1000/48 μm). All values reported for the locations of sCRACM inputs from different presynaptic regions are in multiples of this number. Note, however, that the actual resolution with which synapses can be localised in the heatmaps is likely to be lower than this, as it is limited by both light scattering in the tissue and by the spread of voltage along stimulated axons, which is determined by the length constant of presynaptic axons. Previous studies have indicated that these factors limit the actual sCRACM resolution to approximately 60 μm (Petreanu et al 2009).

To estimate the proportion of sCRACM input targeting different dendritic domains, the recorded input map for each cell was convolved with the average ttL5 morphology obtained from 11 reconstructed Colgalt2-Cre neurons in V2M. This was done by manually separating the apical tuft, oblique (including the apical trunk), and basal dendrites of the reconstructions in Neurolucida 360 and quantifying the total dendritic length in 10 μm thick sections perpendicular to the main axis of the apical dendrite. The resulting dendrite profiles were then aligned by the soma and averaged. Using this average morphology, at each distance from the soma the proportion of dendrites belonging to each domain was calculated relative to the total dendritic length within that section. For each sCRACM recording, these proportions were calculated using an average morphology that was scaled to the soma-pia distance of the recorded cell. Each pixel of the heatmap for that cell was then multiplied by these dendritic proportions to obtain the proportion of evoked current assigned to each dendritic domain. Averaging the morphological reconstructions was necessary because only a small fraction of the recorded neurons was fully reconstructed, and we were thus unable to allocate sCRACM measures to specific dendritic domains on a cell-by-cell basis. Notably, because of variation in apical dendrite length between different ttL5 neurons, the choice of soma alignment before averaging the morphologies resulted in the average morphology profile having a graded rather than sharp cutoff at the pia. The peak dendrite density near the pia thus appears smaller than it would in a pia-aligned average and is not related to any loss of dendrites in the reconstructed neurons. This choice was made because the distribution of tuft dendrites had relatively little overlap with the other dendritic domains, resulting in a more accurate assignment of sCRACM inputs to each domain. Had we aligned the morphologies by the pia instead, the resulting distributions for basal and oblique dendrites would have far greater overlap, resulting in lower accuracy in the dendritic domain classification.

The average morphology profile was also used to quantify the expected input profile on the basis of axon and dendrite densities at each distance from the pia. In this case, however, the same reconstructed morphologies were instead aligned by the apical tuft before averaging, in order to make the predicted input more comparable to the pia-aligned sCRACM maps. As with the sCRACM input, pia-alignment resulted in greater definition at the pia at the cost of reduced resolution near the average soma location. Following alignment, both the axon and dendrite distributions were normalized to the peak of each curve and multiplied, resulting in large values for expected input at locations containing both axons and dendrites.

## Acknowledgments

We thank Troy Margrie and Molly Strom for viral constructs; Rob Campbell and Charlie Rousseau for help with data acquisition and analysis of rabies tracing experiments, and Joe Brock for help with illustrations. We are grateful to Florencia Iacaruso and Zoltán Kisvárday for helpful comments on the manuscript.

## Competing interests

The authors declare that no competing interests exist.

## Supplementary material

**Figure S1.**
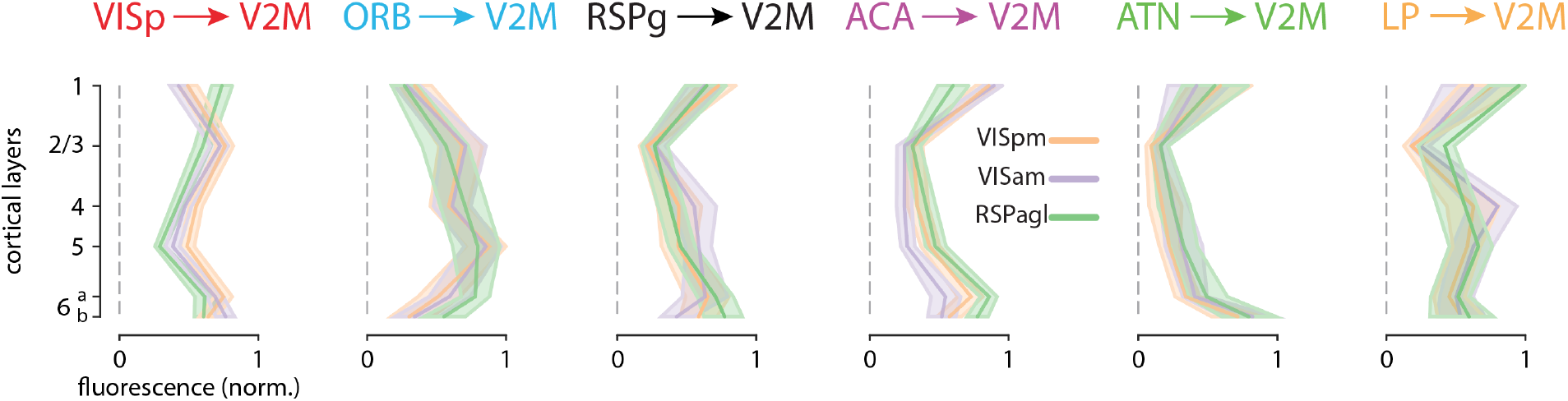
Comparison of axonal projection profiles. Layer-wise axonal projection profiles to VISpm, VISam and RSPagl. Data from the Allen Mouse Brain Connectivity Atlas.

**Figure S2.**
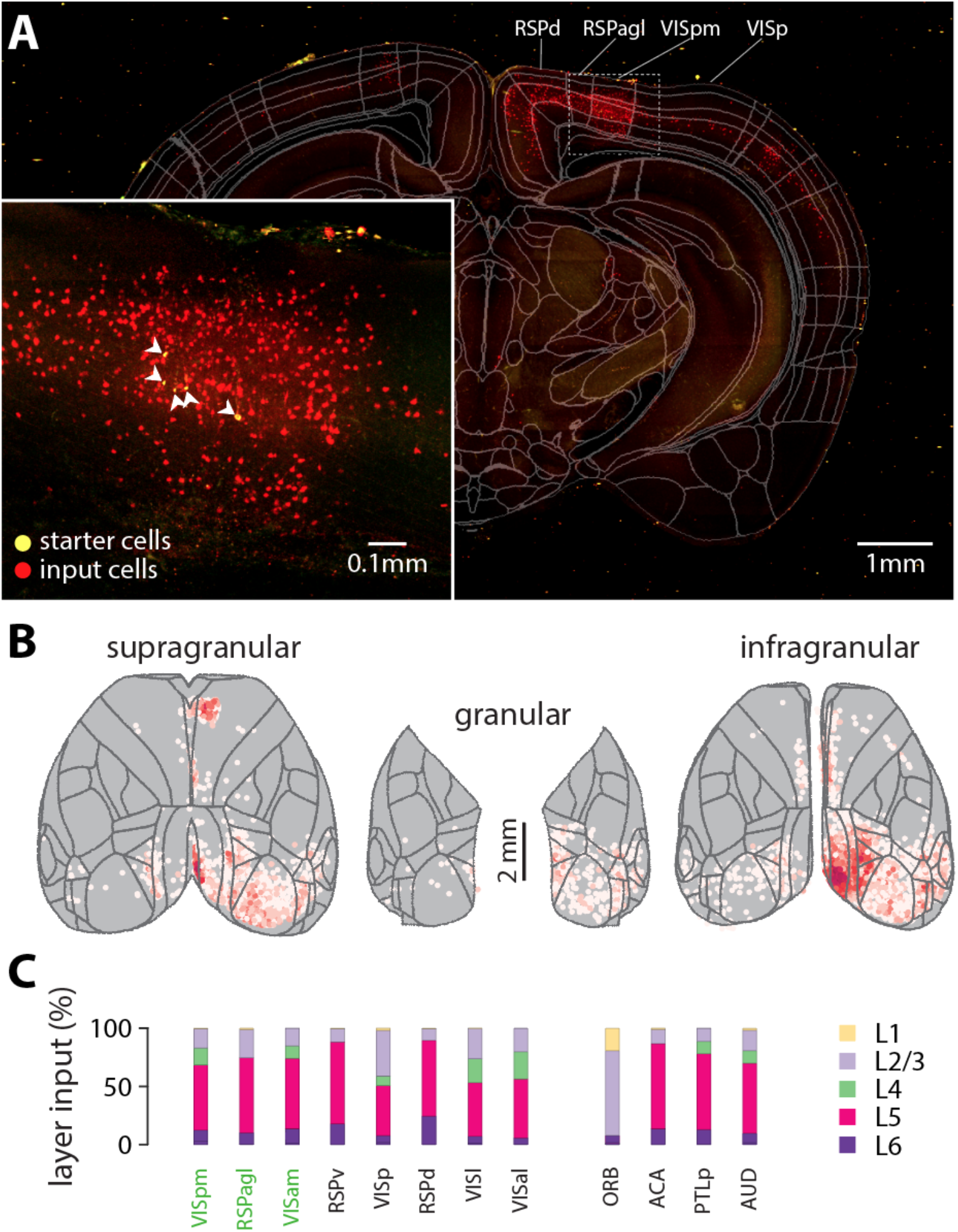
**A.** Maximum intensity projection of a 200 μm thick coronal slab containing some of the starter neurons. The propagated CCFv3 atlas outlines are overlayed. Inset shows starter area with higher magnification. **B.** Input cell density maps across supragranular, granular and infragranular layers. A-B Same experiment and scales as Figure 1. **C.** Distribution of input cells across cortical layers. Average of 3 experiments.

**Figure S3.**
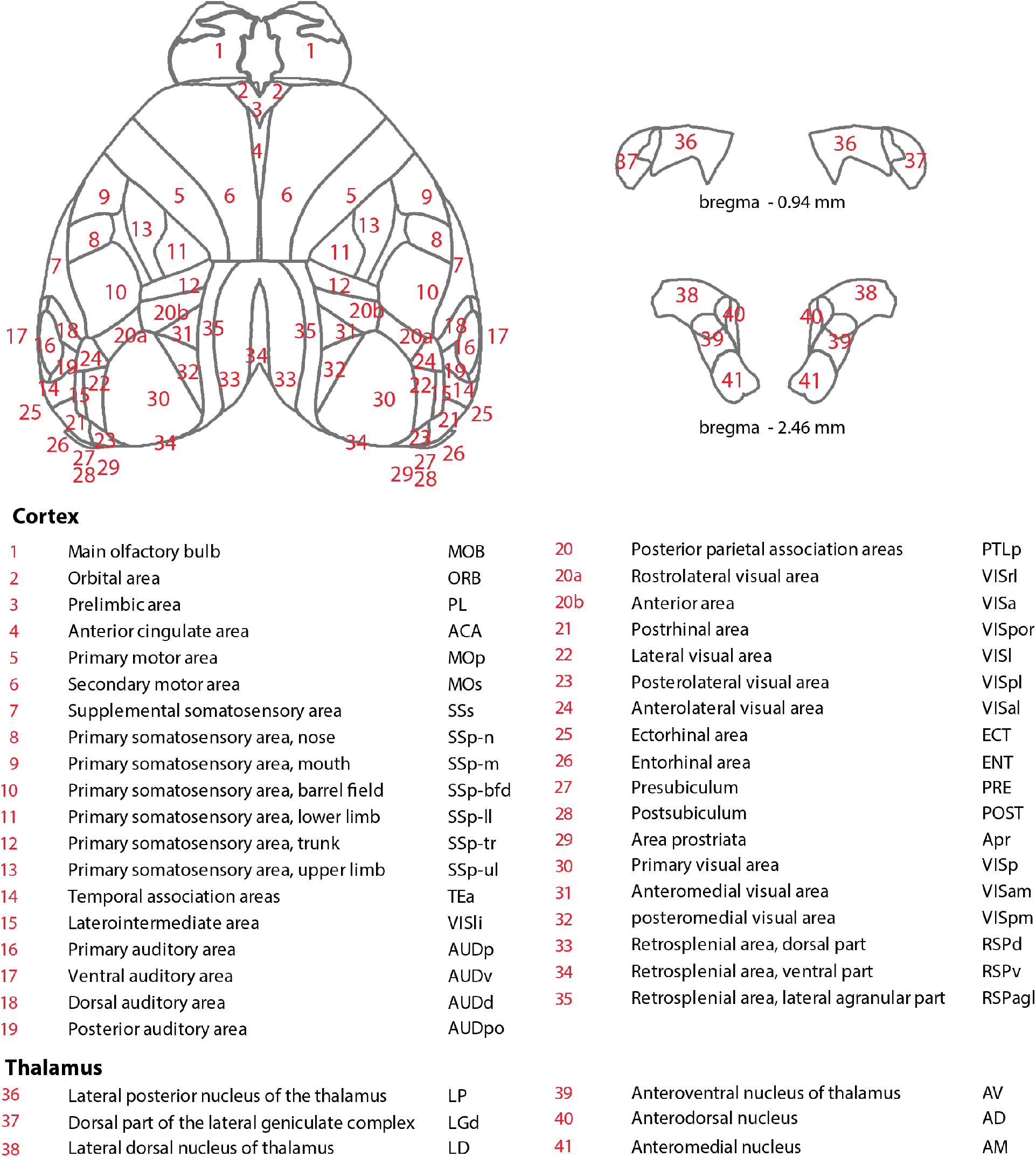
Brain segmentation and nomenclature according to the Allen CCFv3.

**Figure S4.**
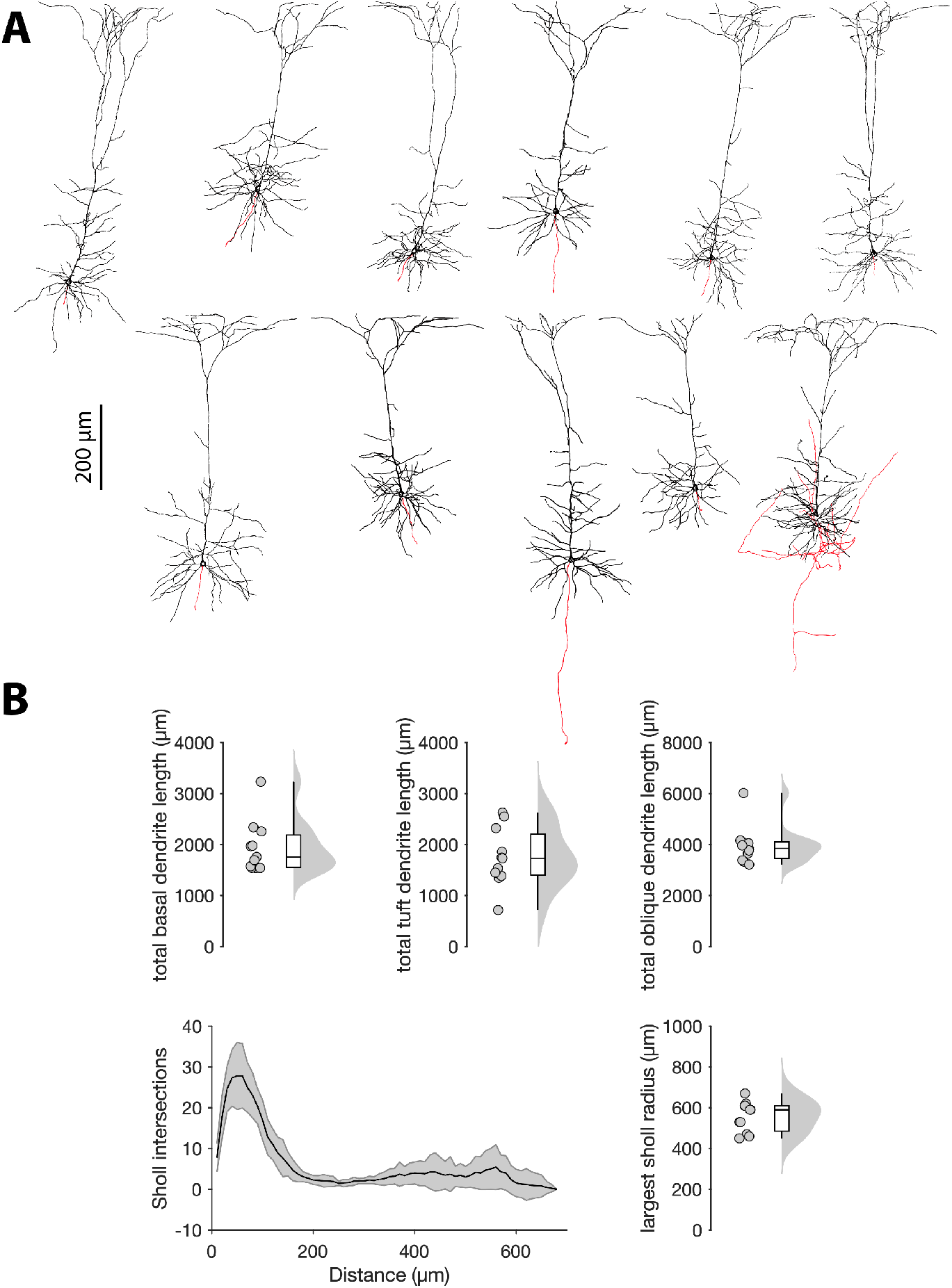
**A.** Reconstructed morphologies of 11 Colgalt-2 neurons. Black: dendrites; red: axons. **B.** Quantitative descriptive measures of dendritic morphology.

**Figure S5.**
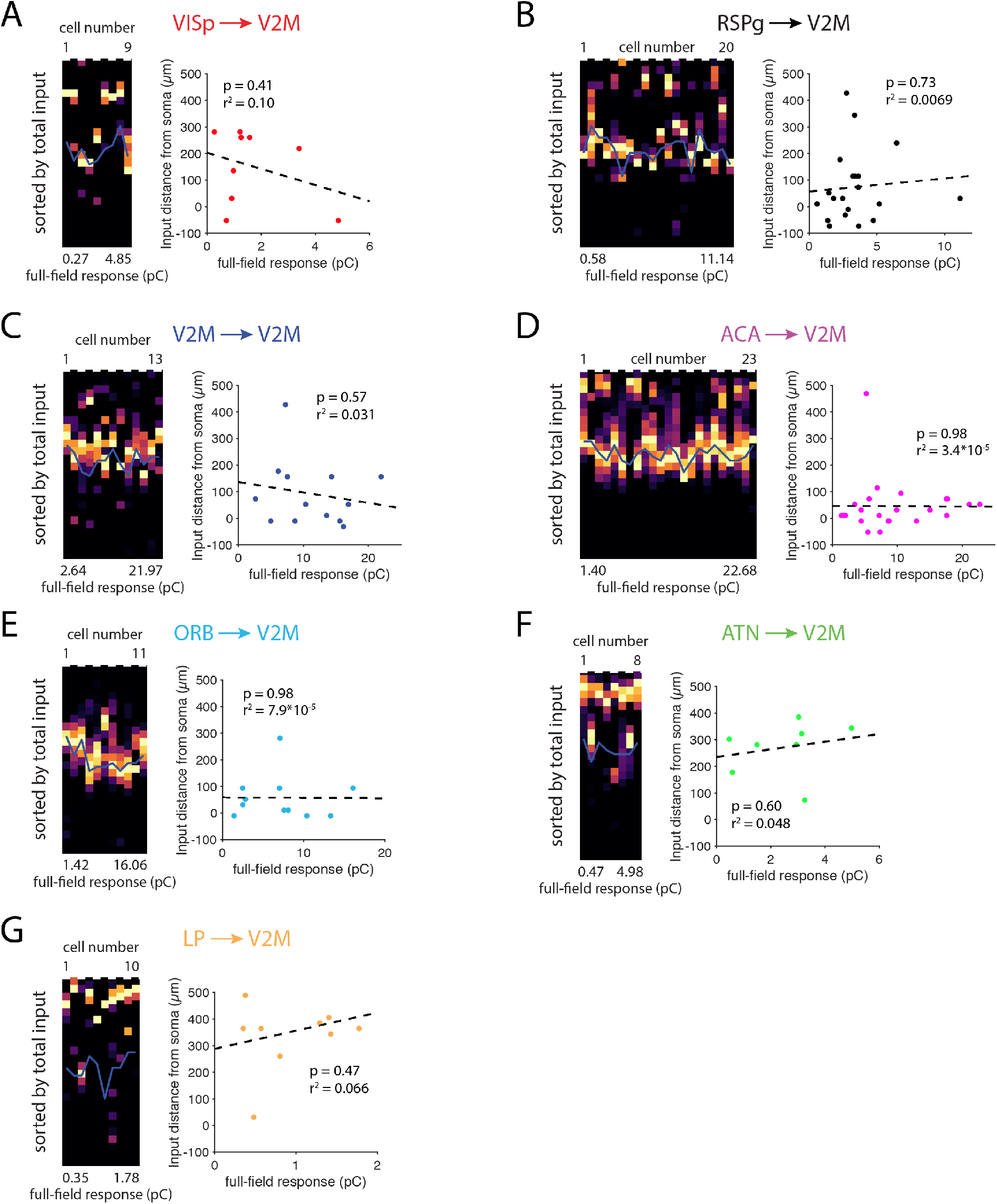
Input location does not correlate with input strength. **A.** Horizontal projections of VISp input to individual cells sorted by full-field stimulation response. Right: Location of the largest input peak versus full-field response. Dashed line is a linear fit. **B-G.** Same as in A, but for RSPg, V2M, ACA, ORB, ATN and LP input.

**Figure S6.**
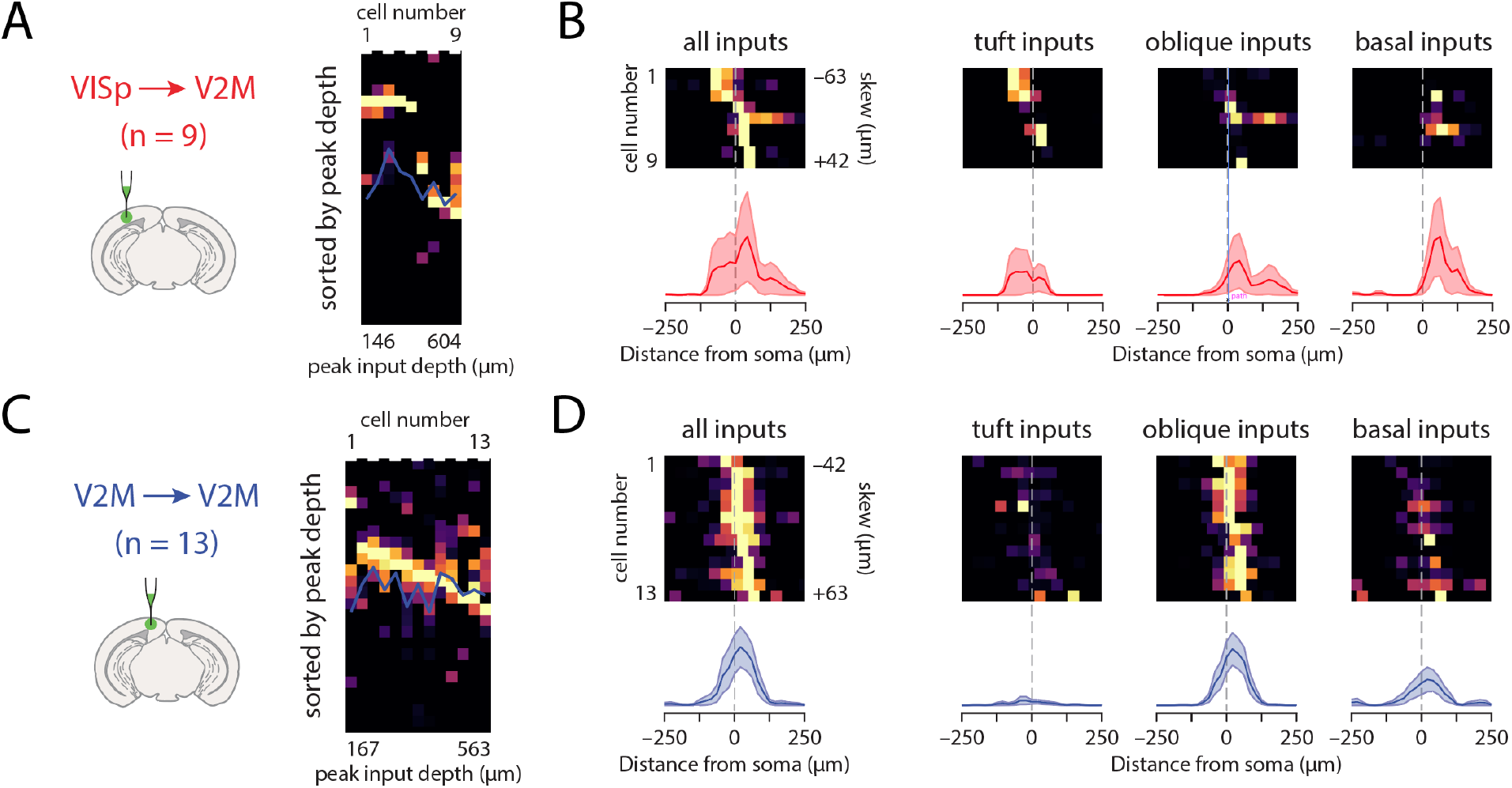
Detailed data analysis for VISp and V2M. **A, C.** Vertical projections of individual input maps sorted by the location of the peak input. **B, D.** Horizontal projections of individual input maps and their average for all inputs, and for domain-separated inputs.

**Figure S7.**
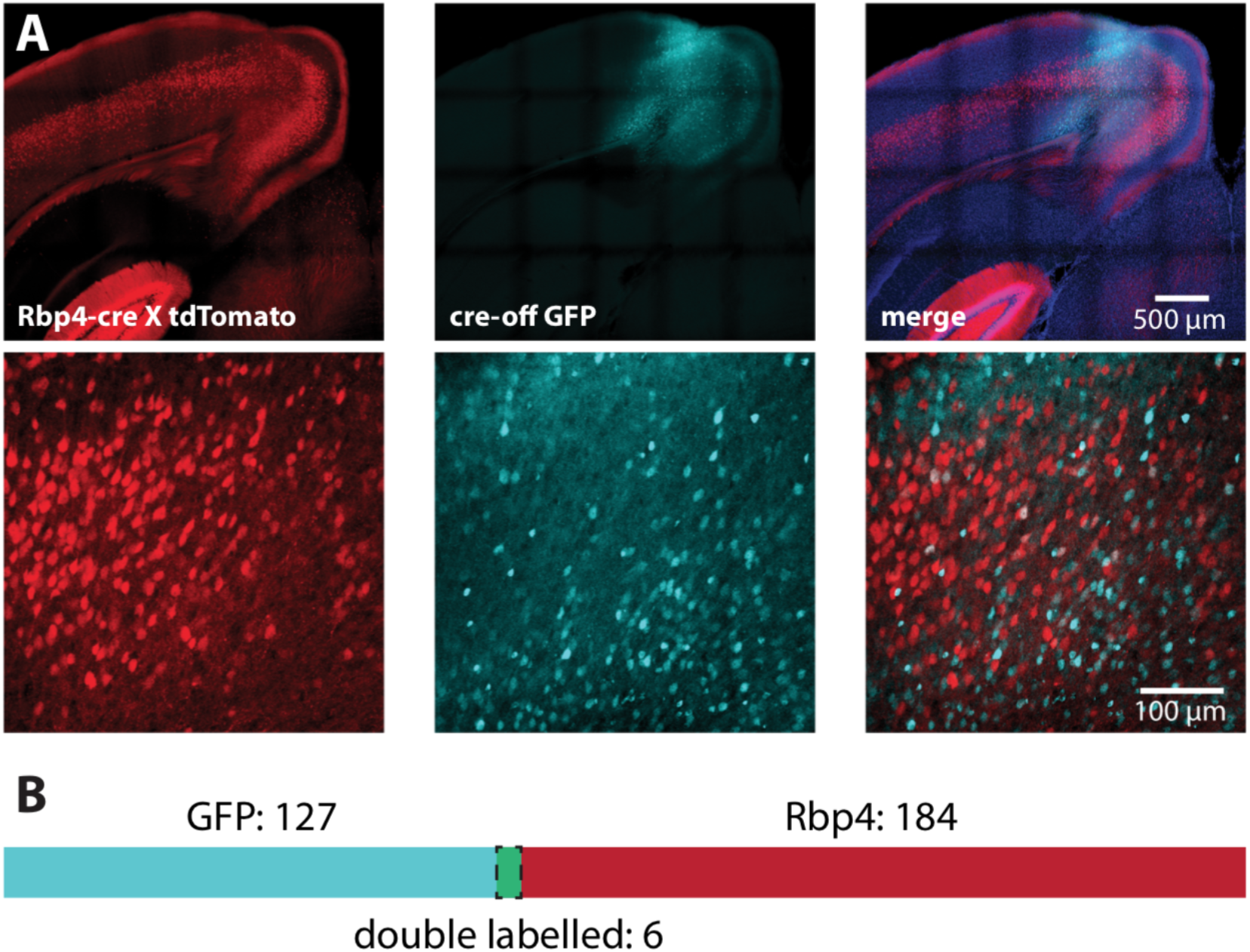
Efficacy of Cre-off virus. **A.** Example confocal images showing cre expression in L5 pyramidal neurons (red), GFP expression following cre-off virus injection (cyan) and overlap (blue = DAPI) on two magnifications. **B.** Quantification of overlap between cre and GFP expression.

**Figure S8.**
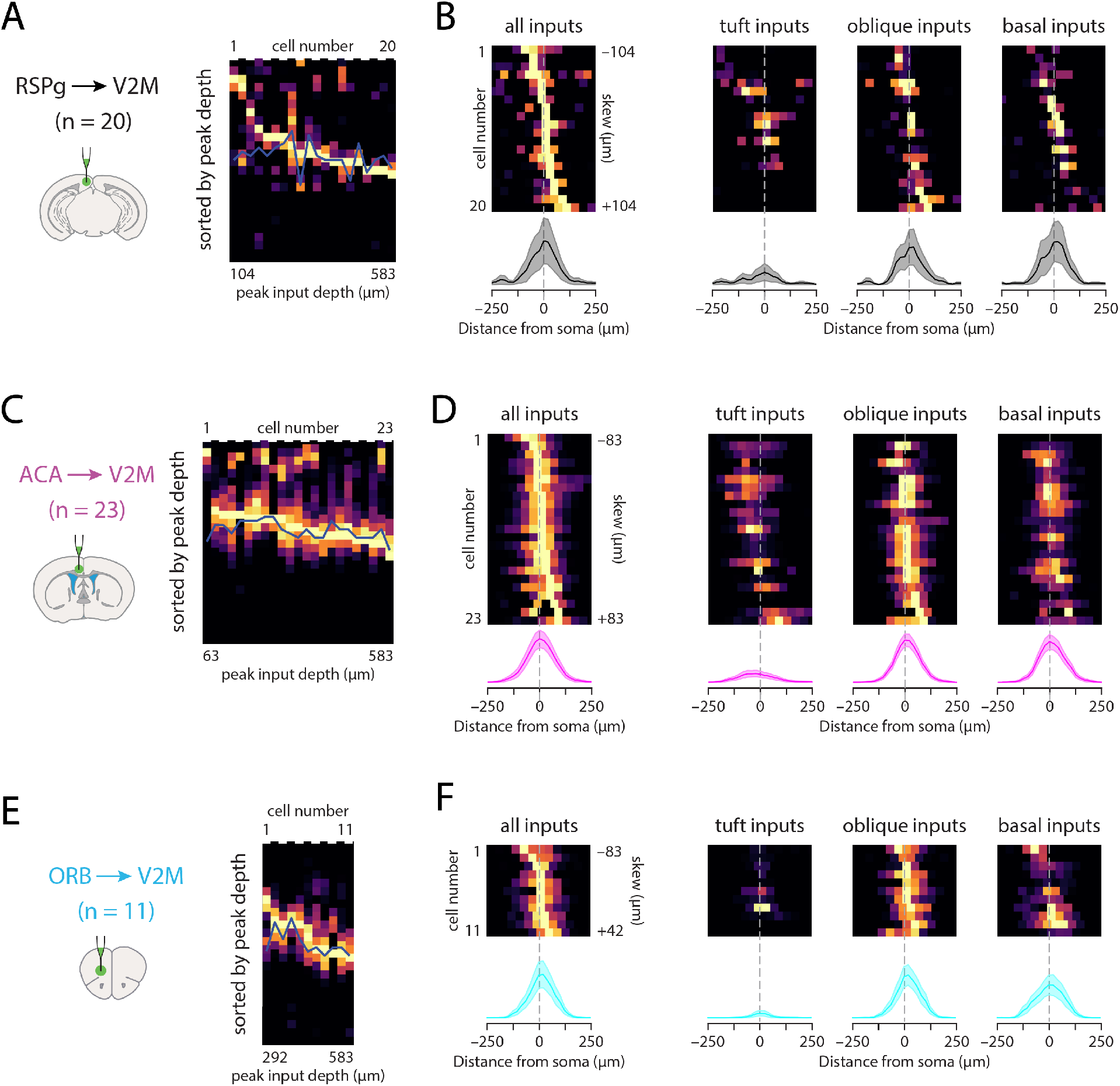
Detailed data analysis for RSPg, ACA and ORB. **A, C, E.** Vertical projections of individual input maps sorted by the location of the peak input. **B, D, F.** Horizontal projections of individual input maps and their average for all inputs, and for domain-separated inputs.

**Figure S9.**
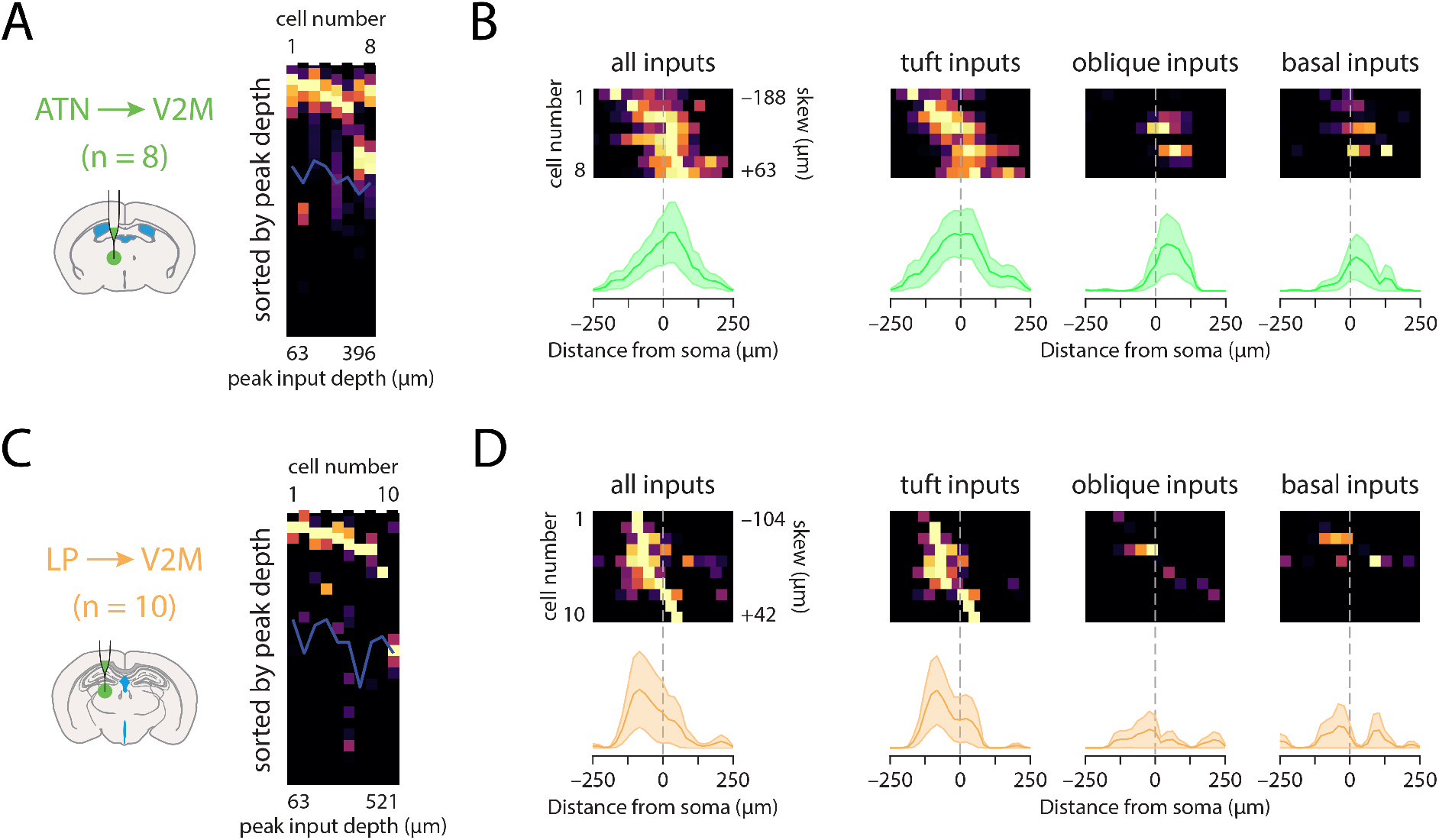
Detailed data analysis for ATN and LP. **A, C.** Vertical projections of individual input maps sorted by the location of the peak input. **B, D.** Horizontal projections of individual input maps and their average for all inputs, and for domain-separated inputs

**Supplementary table 1.** Results of rabies tracing experiments.

| Number of input cells |  |  |  |  |  |  |  |  |  |  |  |
| --- | --- | --- | --- | --- | --- | --- | --- | --- | --- | --- | --- |
| thalamus | average | sd | sample 1 | sample 2 | sample 3 |  |  |  |  |  |  |
| LP | 1191 | 191 | 1409 | 1052 | 1111 |  |  |  |  |  |  |
| ATN | 1001 | 417 | 1465 | 657 | 882 |  |  |  |  |  |  |
| LGd | 194 | 174 | 366 | 198 | 19 |  |  |  |  |  |  |
|  |  |  |  |  |  | Layerwise contribution |  |  |  |  |  |
| proximal cortex |  |  |  |  |  | L1 | L2/3 | L4 | L5 | L6a | L6b |
| VISp | 1245 | 282 | 954 | 1262 | 1518 | 1.83% | 39.36% | 8.10% | 42.97% | 5.97% | 1.77% |
| VISpm | 842 | 264 | 687 | 692 | 1146 | 0.28% | 16.63% | 14.62% | 55.98% | 9.68% | 2.80% |
| VISam | 418 | 227 | 234 | 348 | 672 | 0.00% | 15.12% | 10.75% | 60.37% | 12.43% | 1.32% |
| VISI | 334 | 144 | 170 | 392 | 439 | 0.17% | 25.97% | 20.55% | 46.06% | 5.92% | 1.33% |
| VISal | 118 | 65 | 46 | 174 | 133 | 0.00% | 20.08% | 23.55% | 50.62% | 5.36% | 0.40% |
| RSPv | 1825 | 844 | 2705 | 1022 | 1747 | 0.29% | 11.40% | n/a | 70.43% | 17.87% | 0.01% |
| RSPd | 1038 | 314 | 1256 | 678 | 1180 | 0.29% | 9.97% | n/a | 65.15% | 24.45% | 0.14% |
| RSPagl | 785 | 353 | 967 | 378 | 1010 | 1.14% | 24.25% | n/a | 64.46% | 9.96% | 0.19% |
| distal cortex |  |  |  |  |  |  |  |  |  |  |  |
| ORB | 305 | 117 | 181 | 320 | 414 | 19.25% | 72.86% | n/a | 6.22% | 1.67% | 0.00% |
| ACA | 252 | 161 | 189 | 131 | 435 | 0.96% | 12.20% | n/a | 73.28% | 13.56% | 0.00% |
| PTLp | 221 | 87 | 120 | 266 | 276 | 0.00% | 11.16% | 10.62% | 65.23% | 12.60% | 0.39% |
| AUD | 177 | 28 | 145 | 195 | 192 | 1.58% | 17.60% | 10.83% | 60.17% | 8.18% | 1.64% |

**Supplementary table 2.** Results of all sCRACM experiments.

| input area | parameter | basal | oblique | tuft* |  |  |
| --- | --- | --- | --- | --- | --- | --- |
| **VISp**<br>N = 6<br>n = 9 | peak location ($\mu\text{m}$) | -62.5 | 104.17 | 187.5 | total input (pC) | 0.93 |
|  | input proportion (%) | 26% | 33% | 42% | total input SEM | 0.11 |
| | proportional input (pC) | 0.24 | 0.30 | 0.39 | soma depth ($\mu\text{m}$) | 507 ± 22 |
| | horizontal bias ($\mu\text{m}$) ** | 62.5 | 41.67 | -20.83 | cells with peak in tuft | 5/9 |
| **V2M**<br>N = 4<br>n = 13 | peak location ($\mu\text{m}$) | -41.67 | 83.33 | 166.67 | total input (pC) | 11.24 |
|  | input proportion (%) | 24% | 62% | 14% | total input SEM | 1.56 |
| | proportional input (pC) | 2.68 | 6.96 | 1.60 | soma depth ($\mu\text{m}$) | 498 ± 15 |
| | horizontal bias ($\mu\text{m}$) ** | 20.83 | 20.83 | -20.83 | cells with peak in tuft | 1/13 |
| **RSPg**<br>N = 9<br>n = 20 | peak location ($\mu\text{m}$) | -41.67 | 41.67 | 125 | total input (pC) | 3.40 |
|  | input proportion (%) | 30% | 40% | 30% | total input SEM | 0.51 |
| | proportional input (pC) | 1.04 | 1.36 | 1.01 | soma depth ($\mu\text{m}$) | 503 ± 15 |
| | horizontal bias ($\mu\text{m}$) ** | 20.83 | 20.83 | 0 | cells with peak in tuft | 2/20 |
| **ACA**<br>N = 5<br>n = 23 | peak location ($\mu\text{m}$) | -41.67 | 41.67 | 83.33 | total input (pC) | 9.46 |
|  | input proportion (%) | 30% | 45% | 25% | total input SEM | 1.32 |
| | proportional input (pC) | 2.83 | 4.22 | 2.41 | soma depth ($\mu\text{m}$) | 464 ± 9 |
| | horizontal bias ($\mu\text{m}$) ** | 0 | 0 | -20.83 | cells with peak in tuft | 1/23 |
| **ORB**<br>N = 3<br>n = 11 | peak location ($\mu\text{m}$) | -41.67 | 20.83 | N/A | total input (pC) | 7.16 |
|  | input proportion (%) | 35% | 57% | 9% | total input SEM | 1.43 |
| | proportional input (pC) | 2.47 | 4.05 | 0.63 | soma depth ($\mu\text{m}$) | 521 ± 19 |
| | horizontal bias ($\mu\text{m}$) ** | 0 | 20.83 | 0 | cells with peak in tuft | 0/11 |
| **ATN**<br>N = 3<br>n = 8 | peak location ($\mu\text{m}$) | -104.17 | 104.17 | 104.17 | total input (pC) | 2.48 |
|  | input proportion (%) | 8% | 17% | 75% | total input SEM | 0.54 |
| | proportional input (pC) | 0.21 | 0.42 | 1.86 | soma depth ($\mu\text{m}$) | 435 ± 15 |
| | horizontal bias ($\mu\text{m}$) ** | 20.83 | 41.67 | 20.83 | cells with peak in tuft | 6/8 |
| **LP**<br>N = 4<br>n = 10 | peak location ($\mu\text{m}$) | -41.67 | 20.83 | 62.5 | total input (pC) | 0.97 |
|  | input proportion (%) | 10% | 15% | 75% | total input SEM | 0.16 |
| | proportional input (pC) | 0.10 | 0.14 | 0.72 | soma depth ($\mu\text{m}$) | 500 ± 23 |
| | horizontal bias ($\mu\text{m}$) ** | -41.67 | -20.83 | -83.33 | cells with peak in tuft | 9/10 |
|  | **total proportional input** | **27%** | **49%** | **24%** | **sum total input (pC)** | **35.65** |

\*tuft measurements from pia, basal and oblique from soma
\*\* negative means lateral, positive medial

**Supplementary table 3.** Stereotaxic coordinates and volumes of viral injections.

| Area | Distance from bregma (mm) | Mediolateral distance (mm) | Depth from pia (mm) | Number & volume of injections |
| --- | --- | --- | --- | --- |
| VISp | [-3.5 : -2.8] | [1.8 : 2.7] | [0.5 : 0.6] | 3 x 100 nL |
| RSPg | [-3.2 : -2.7] | 0.5 | [0.5 : 0.7] | 2 x 100 nL |
| V2m | [-3.2 : -2.7] | 1 | [0.2 : 0.5] | 2 x 100 nL |
| ACA | [0 : 1] | 0.5 | [1.2 : 1.5] | 3 x 100 nL |
| ORB | [2 : 2.8] | 1 | [1.5 : 2.3] | 3 x 100 nL |
| ATN | [-0.5 : -1.2] | [0.5 : 0.7] | [3.2 : 3.3] | 2 x 100 nL |
| LP | [-2.5 : -1.7] | [1 : 1.5] | [2.4 : 2.6] | 3 x 100 nL |

## Notes

### Competing Interest Statement

The authors have declared no competing interest.

